# One particle per residue is sufficient to describe all-atom protein structures

**DOI:** 10.1101/2023.05.22.541652

**Authors:** Lim Heo, Michael Feig

**Affiliations:** Department of Biochemistry and Molecular Biology, Michigan State University, East Lansing, MI 48824, USA

**Keywords:** Biological Sciences, Biophysics and Computational Biology

## Abstract

Atomistic resolution is considered the standard for high-resolution biomolecular structures, but coarse-grained models are often necessary to reflect limited experimental resolution or to achieve feasibility in computational studies. It is generally assumed that reduced representations involve a loss of detail, accuracy, and transferability. This study explores the use of advanced machine-learning networks to learn from known structures of proteins how to reconstruct atomistic models from reduced representations to assess how much information is lost when the vast knowledge about protein structures is taken into account. The main finding is that highly accurate and stereochemically realistic all-atom structures can be recovered with minimal loss of information from just a single bead per amino acid residue, especially when placed at the side chain center of mass. High-accuracy reconstructions with better than 1 Å heavy atom root-mean square deviations are still possible when only Cα coordinates are used as input. This suggests that lower-resolution representations are essentially sufficient to represent protein structures when combined with a machine-learning framework that encodes knowledge from known structures. Practical applications of this high-accuracy reconstruction scheme are illustrated for adding atomistic detail to low-resolution structures from experiment or coarse-grained models generated from computational modeling. Moreover, a rapid, deterministic all-atom reconstruction scheme allows the implementation of an efficient multi-scale framework. As a demonstration, the rapid refinement of accurate models against cryoEM densities is shown where sampling at the coarse-grained level is guided by map correlation functions applied at the atomistic level. With this approach, the accuracy of standard all-atom simulation based refinement schemes can be matched at a fraction of the computational cost.

**STATEMENT OF SIGNIFICANCE:** The fundamental insight of this work is that atomistic detail of proteins can be recovered with minimal loss of information from highly reduced representations with just a single bead per amino acid residue. This is possible by encoding the existing knowledge about protein structures in a machine-learning model. This suggests that it is not strictly necessary to resolve structures in atomistic detail in experiments, computational modeling, or the generation of protein conformations via neural networks since atomistic details can inferred quickly via the neural network. This increases the relevance of experimental structures obtained at lower resolutions and broadens the impact of coarse-grained modeling.

## INTRODUCTION

Proteins play central roles in biological processes, and their behavior is often studied at the molecular level to understand biological function. An atomistic level of resolution is the gold standard for experiments and computation alike. Experimental methods such as X-ray crystallography (1, 2), nuclear magnetic resonance (NMR) (3), and cryogenic electron microscopy (cryo-EM) (4) can resolve structures in atomistic detail, but achieving such high resolution requires significant effort (5, 6). Computational modeling and simulations also typically require all-atom representations of the protein to achieve maximum accuracy and to gain detailed mechanistic insights (7, 8). Atomistic modeling remains computationally expensive, though, limiting practical applications (9), even with the latest high-performance computing platforms and simulation accelerators. (*e.g.*, Anton or CUDA) (10). Similarly, it is also very demanding to train machine-learning based methods for directly predicting atomistic models (11-13) and conformational ensembles (14, 15).

Coarse-graining (CG) of protein structures is a common strategy to overcome the various challenges (7). When interpreting experimental data, reduced representations may be a natural fit to match lower experimental resolutions. In computational applications, CG models greatly increase efficiency by reducing the number of particles. CG representations may range from single beads per protein (16, 17) to residue-based models (18, 19) and multiple sites per amino acid residue (20-24). Lower-resolution models of experimental data often default to Cα traces. In computational applications, the choice of resolution may depend on the questions that are being investigated as model accuracy and transferability depend on the degree of coarse-graining (7, 25).

To recover atomistic information from CG models, all-atom reconstruction algorithms have been developed with different strategies depending on the CG representation. For a united-atom model, which omits only hydrogen atoms, missing hydrogens can be placed using their pre-defined local geometries (26). An all-atom structure can still be generated relatively accurately and quickly from higher-resolution CG models such as PRIMO or MARTINI, based on geometry-based reconstruction rules (21, 27). The reconstruction of all-atom structures from coarser representations such as Cα-traces is more complex. Methods such as PULCHRA (28) and REMO (29) convert Cα-traces to all-atom structures by first rebuilding the backbone atoms before predicting side chain orientations. These methods typically rely on pre-defined backbone fragment libraries, side chain rotamer libraries (30), or other empirical information derived from known structures. Extensive optimization is then often required to avoid clashes and find energetically optimal structures (31). Nevertheless, the resulting reconstructions may retain significant deviations from correct all-atom structures when only Cα atoms are available as input. The reconstructions may also vary from one run to another if they depend on stochastic optimization techniques. The relatively poor accuracy when reconstructing atomistic detail from lower resolutions has limited the full interpretation of experimental data that does not directly provide atomistic details and hindered effective implementations of multi-scale sampling methods that are both efficient and thermodynamically consistent with sampling at all-atom levels (8, 32, 33).

In the meantime, the recent success of accurate structure prediction via machine learning methods (11-13) has demonstrated that sufficiently deep neural network models can effectively learn from the large amount of known structures how to generate atomistic models just from amino acid sequences. This suggests that it should also be possible using similar approaches to reconstruct atomistic detail at high resolution if lower-resolution structural information is available as additional input.

Inspired by AlphaFold2 (11), we trained an SE(3)-equivariant graph neural network model for reconstructing all-atom detail from lower representations. Like AF2, the model utilizes rigid-body blocks for generating 3D structures from predicted features, but the model was extended to better describe hydrogen atoms and secondary structure-dependencies. The network learned structural features of backbone and sidechain atoms from known protein conformations, but also incorporates physical constraints necessary to produce realistic all-atom structures. The model is applicable to a range of CG models such as Cα-traces, traces of residue-center-of-mass model, and MARTINI models (20). It provides all-atom reconstructions at much higher accuracy than with previous methods, better than 1 Å for heavy atoms from only Cα atoms and better than 0.5 Å with a single site at reside centers of mass. From a more general perspective this suggests that atomistic details of proteins at a resolution close to experimental accuracy can be captured essentially with a single site per residue if the current knowledge of protein structures is taken advantage of via machine learning.

The all-atom reconstruction via the machine-learning framework is fast and deterministic, and since gradients are available via back propagation, it is straightforward to map energies and constraints at the all-atom level directly to the CG representation. It is therefore possible to sample a residue-based model guided directly by all-atom forces via back-mapping through the all-atom reconstruction network. As a proof-of-principle we demonstrate practical value in the rapid refinement of all-atom coordinates against intermediate- and low-resolution cryo-EM densities. The protocol achieves comparable accuracy to traditional all-atom simulation-based approaches but with much reduced computational effort.

## RESULTS AND DISCUSSION

### Accurate reconstruction of all-atom structures from coarse-grained representations

We trained SE(3)-equivariant machine learning models, called cg2all, to reconstruct all-atom structures of proteins from CG representations (*cf.* Methods section and *SI Appendix*). The network architecture is shown in **Fig. 1**. Model variations with different hyperparameters were explored to determine the optimal model architecture (*SI Appendix*). An ablation study was further carried out to determine optimal input features and loss function components (*SI Appendix*). Only results with the optimized architecture are described subsequently.

**Figure 1.**
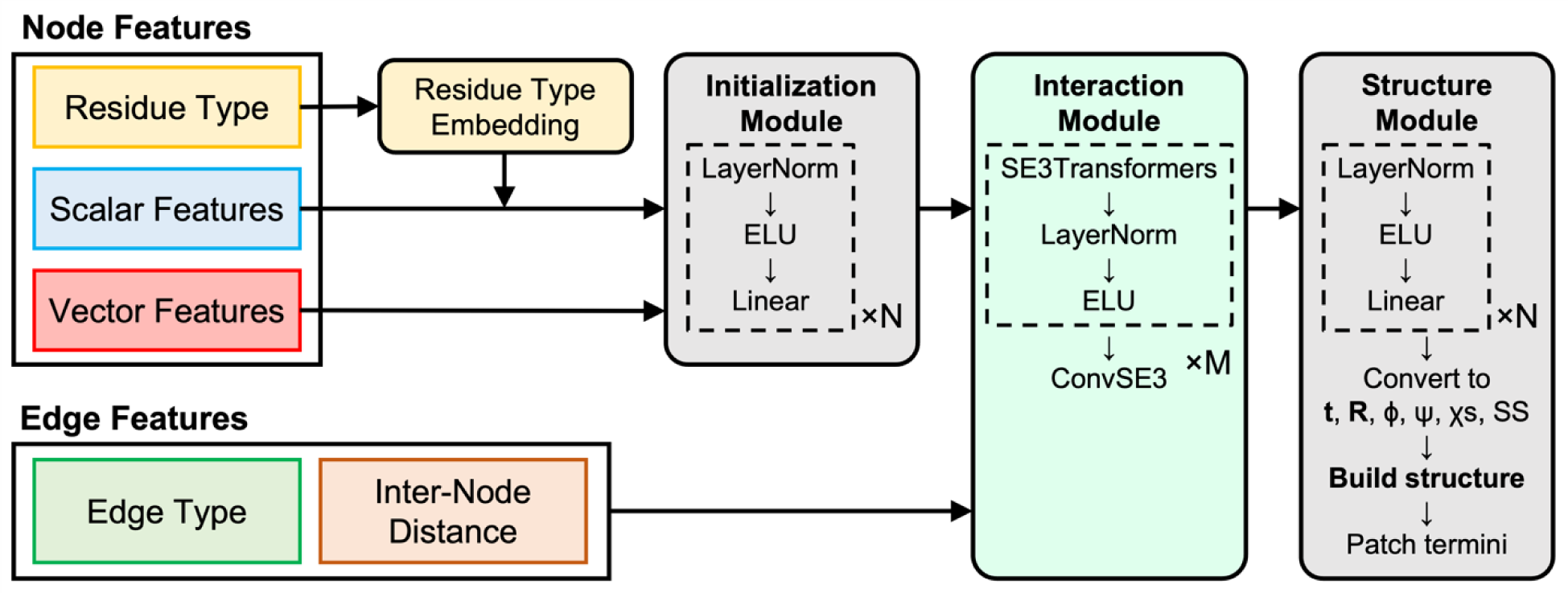
Architecture of coarse-grained to all-atom structure conversion model.

Models were generated for the reconstruction from Cα atoms, from all backbone atoms, from a single particle at the center of mass of an amino acid residue, from the MARTINI model with several beads per residue (20), and from the higher-resolution CG model PRIMO (21). The generated all-atom models were evaluated on a validation set in terms of root mean square deviations (RMSD) and side chain torsion accuracy with respect to the original reference structures as well as MolProbity (34) scores to check stereochemical quality (**Table 1**). Because the machine-learning model estimates internal parameters that are then used to reconstruct all-atom detail via rigid-body reconstruction (11), using the parameters from the experimental all-atom structures as input for the rigid-body reconstruction provides an upper limit on the accuracy that can be achieved theoretically. In this ideal case, heavy atom RMSDs of 0.16 Å and MolProbity scores of 1.81 are obtained (**Table 1**), both are within experimental accuracy (34, 35). The reconstruction from the highest-resolution CG model, PRIMO, reaches the theoretical maximum accuracy and even lower MolProbity scores are obtained with slightly fewer clashes (**Table 1**). That may be expected since PRIMO was designed to retain maximum information from all-atom representations. However, even with lower-resolution models, it is still possible to recover highly accurate all-atom structures. Reconstruction from MARTINI models resulted only in a slight loss of accuracy (0.31 Å RMSD) and only slightly increased MolProbity scores. Remarkably, even a single bead per residue, located at the center of a residue, still allows highly accurate reconstruction of all-atom details (0.46 Å RMSD) without significant compromise of stereochemical quality. If the CG site is located at the Cα position, as is common in many CG representations, the loss of accuracy is greater with the average heavy-atom RMSD approaching 1 Å RMSD. The reason is that it becomes more difficult to accurately position side chains if only backbone atoms are given, especially side chains on the surface that are inherently free to sample different rotamer states (**Fig. 2A**). On the other hand, residue center-of-mass models contain information about the location of the side chain position and therefore side chains can be placed accurately, even on the surface (**Fig. 2B**). A more detailed analysis on the models reconstructed from Cα-traces and residue center-of-mass models shows that backbone and side chain torsion angles are closely matched (*SI Appendix* **Fig. S1**). The backbone angles (Cα-C-N and C-N-Cα) are also closely matched (*SI Appendix* **Fig. S2**), but the peptide bond (C-N) showed a somewhat larger standard deviation of 0.027 Å around the average distance of 1.322 Å in the reconstructed all-atom structures from Cα-traces (or 1.330 ± 0.048 Å from residue center-of-mass models) compared to a standard deviation of 0.008 Å around an average of 1.331 Å in the experimental structures. The greater variation in the only flexible backbone bond distance in the rigid-body reconstruction procedure likely compensated for keeping all other bonds rigid. However, one should also note that all but the very highest resolution experimental structures are solved using molecular modeling programs that bias experimental structures towards expected bond lengths (36). This likely results in apparently reduced variations of such bonds. Finally, *cis*-peptide ω torsion angles were not produced for non-pre-proline residues (*SI Appendix* **Fig. S3**).

**Figure 2.**
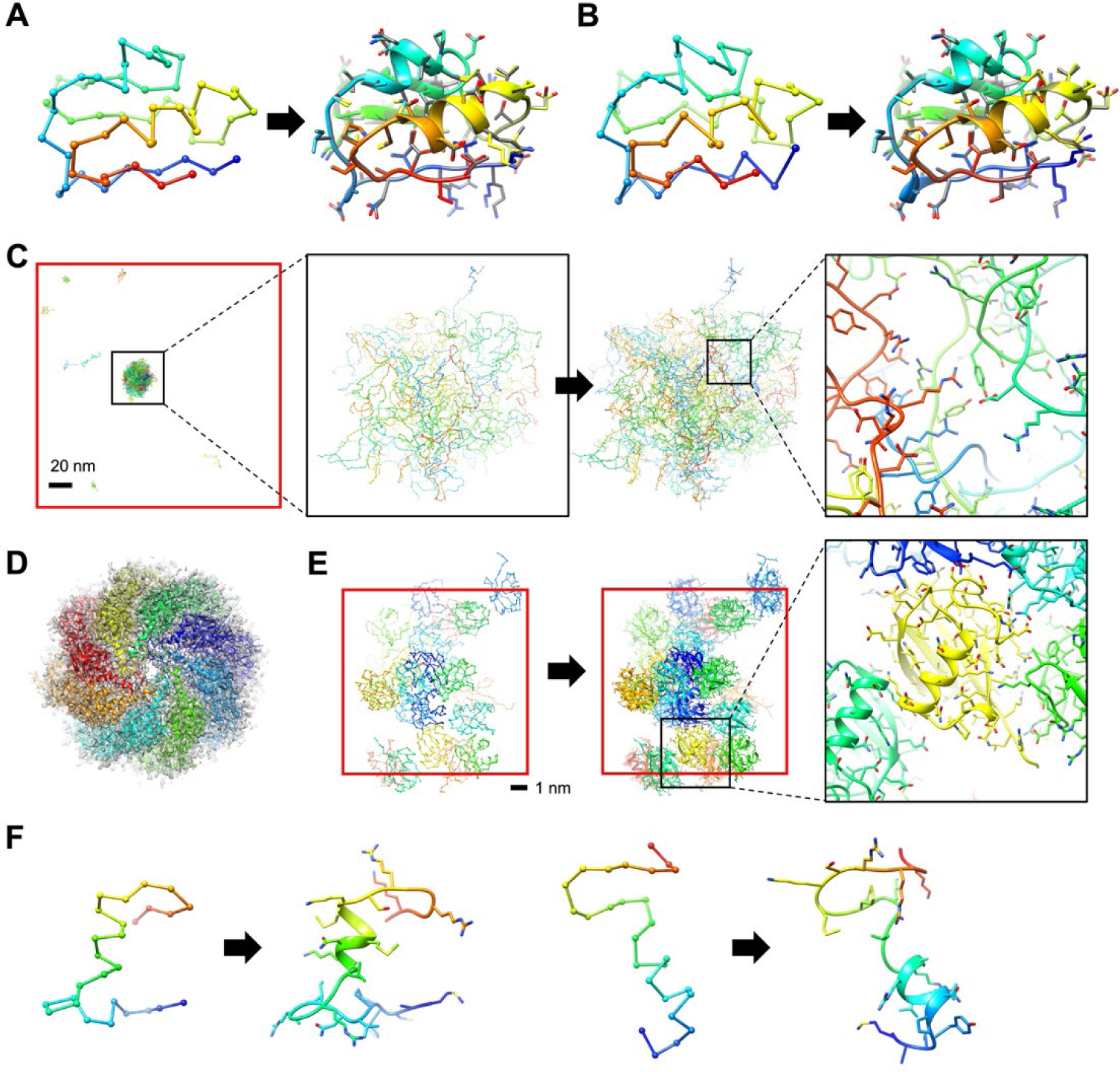
Examples of conversion from CG models to all-atom models. An X-ray crystal structure (PDB ID: 1vjw (37)) recovery from its Cα-trace (A) and residue center-of-mass model (B). The recovered structure is shown as rainbow-colored cartoon representation (from blue to red for N- to C-termini), while its X-ray crystal structure is shown in grey. (C) Conversion of a COCOMO (18) simulation snapshot from a simulation of liquid-liquid phase separation (LLPS) of LAF-1 RGG peptides at 0.042 mM (38). Each peptide consisted of 168 residues, and there were 84 monomers (14,112 residues in total). They are shown in different colors. The phase separated particles at the CG level, and a local region after the conversion are magnified to present detailed structure information (black boxes). The conversion took 33.5 seconds using 16 CPU threads. (D) Building an all-atom model from a medium resolution cryo-EM Cα-trace (PDB ID: 3iyg (39), EMDB ID: 5148, resolution: 4.0 Å). Each chain is depicted in a different color, and the electron density is overlaid as transparent grey voxels. The density map correlation with the all-atom model was 0.723 (*vs.* 0.647 with the Cα-trace). The conversion of the 4,134-residue protein took 11.9 seconds using 16 CPU threads. (E) Conversion of a CG MD simulation trajectory of folded proteins using COCOMO. There are 24 ubiquitins (1,824 residues in total) in the simulation box (shown in red) with a width of 10.76 nm, which resulted in a concentration of 32 mM (274.1 g/L). Each monomer is shown in a different color. A local region after the conversion is zoomed in to show atomistic details of interactions between proteins (black boxes). The conversion of 10,000 frames took 1,774 seconds in total using the “cuda” environment with a batch size of 4 (0.15 s/frame for the forward-pass only). (F) Models generated by idpGAN (14) for an intrinsically disordered protein (UniProt ID: Q9EP54, 27 residues). The conversion of 25,000 models took 64.6 seconds in total using the “cuda” environment with a batch size of 250 (1.3 ms/model for the forward-pass only) Hydrogens were reconstructed but are omitted for clarity.

**Table 1.**
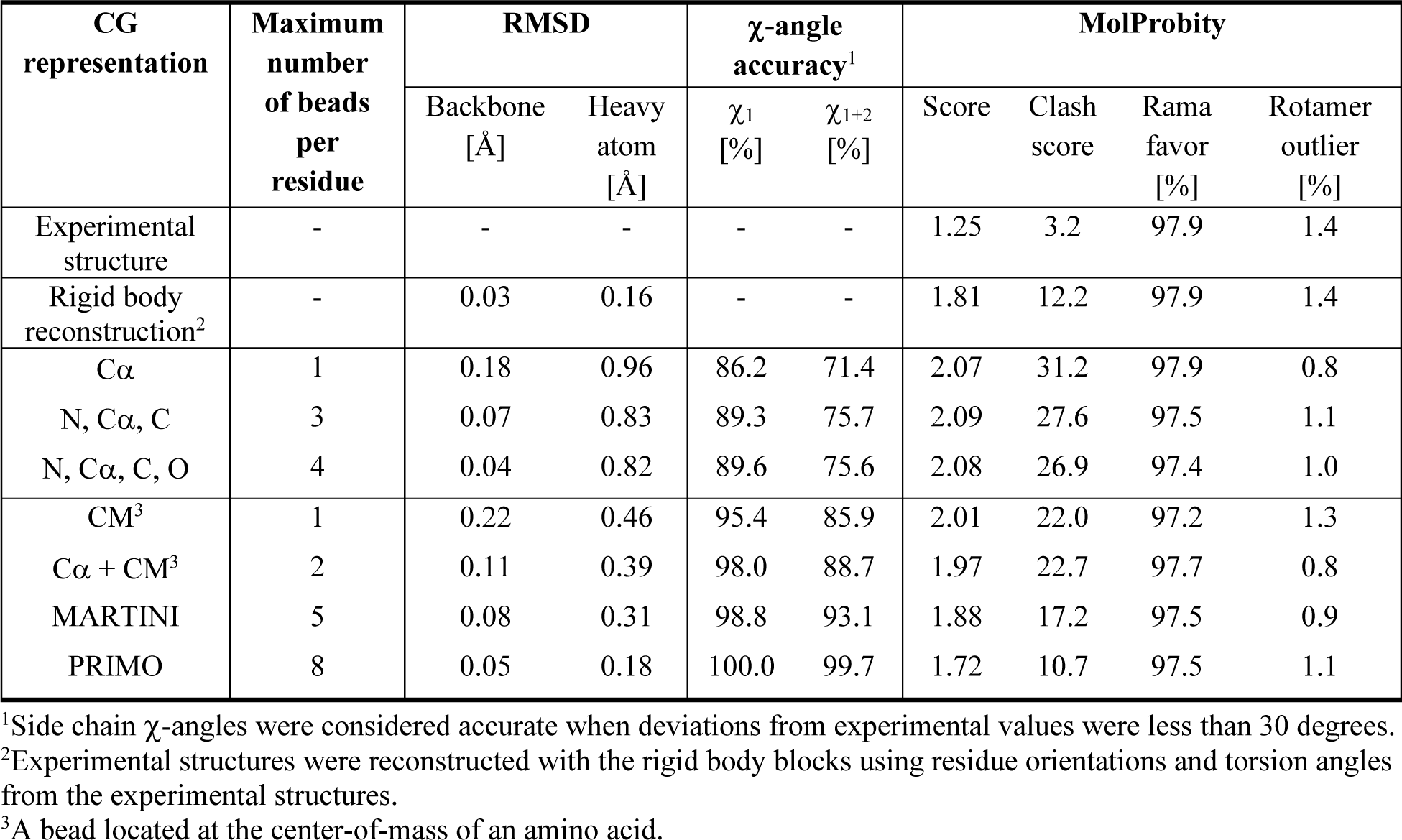
Performance of conversion to all-atom structures from CG models with cg2all.

The machine-learning based all-atom reconstruction via cg2all performed significantly better than previously proposed all-atom reconstruction schemes across all metrics (**Table 2**). No other method achieved significantly better than 1 Å RMSD for heavy atoms, even when reconstructing from the higher-resolution MARTINI model (27). The other older methods also produced structures with significant clashes and higher MolProbity scores, despite energy-guided optimization to avoid clashes. We furthermore tested the widely used rotamer-based method SCRWL4 (30) for placing side chains in combination with backbone atoms generated with cg2all or other methods. Using SCWRL, MolProbity scores were generally improved, even slightly over cg2all, but the accuracy decreased compared to cg2all, especially for MARTINI and center-of-mass based reconstructions. The reason is that SCWRL’s side chain modeling only uses backbone coordinates as input and is based strictly on a rotamer library that, by design, prevents outliers that are occasionally found in experimental structures.

**Table 2.** Comparison of all-atom reconstruction accuracies with different methods.

| Input | Method | RMSD | | χ-angle accuracy<sup>1</sup> | | MolProbity | | | |
| --- | --- | --- | --- | --- | --- | --- | --- | --- | --- |
| | | Backbone<br>[Å] | Heavy<br>atom<br>[Å] | $\chi_1$<br>[%] | $\chi_{1+2}$<br>[%] | Score | Clash<br>score | Rama<br>favor<br>[%] | Rotamer<br>outlier<br>[%] |
| Cα | cg2all<br>w/SCRWL<sup>2</sup> | **0.18** | **0.96**<br>1.06 | **86.2**<br>83.2 | **71.4**<br>70.1 | 2.07<br>**2.00** | 31.2<br>**28.6** | **97.9**<br>**97.9** | 0.8<br>**0.0** |
|  | PULCHRA<sup>3</sup><br>w/SCWRL<sup>2</sup> | 0.47 | 1.58<br>1.36 | 59.2<br>73.0 | 39.6<br>58.4 | 3.79<br>2.90 | 164.4<br>67.4 | 86.4<br>86.4 | 4.9<br>0.1 |
|  | REMO<sup>3,4</sup><br>w/SCWRL<sup>2</sup> | 0.82 | 2.09<br>1.74 | 43.6<br>68.5 | 28.5<br>53.8 | 4.37<br>3.15 | 200.2<br>95.7 | 78.4<br>78.4 | 14.8<br>0.2 |
|  | ModRefiner<sup>3,4</sup> | 0.66 | 1.51 | 71.8 | 55.1 | 2.38 | 56.5 | 97.0 | 0.6 |
|  | MODELLER | 0.98 | 2.12 | 42.8 | 25.4 | 3.63 | 93.3 | 86.4 | 6.0 |
| CM | cg2all<br>w/SCWRL<sup>2</sup> | **0.22** | **0.46**<br>1.00 | **95.4**<br>84.0 | **85.9**<br>70.9 | **2.01**<br>**2.01** | **22.0**<br>25.5 | **97.2**<br>97.2 | 1.3<br>**0.0** |
|  | PULCHRA<sup>3</sup><br>w/SCWRL<sup>2</sup> | 1.08 | 1.91<br>1.84 | 46.6<br>55.3 | 29.0<br>36.3 | 4.49<br>3.51 | 291.6<br>176.2 | 72.9<br>72.9 | 10.5<br>0.2 |
|  | MODELLER<br>(40) | 1.63 | 2.38 | 43.7 | 25.5 | 3.80 | 123.6 | 81.8 | 5.5 |
| MARTINI | cg2all<br>w/SCWRL<sup>2</sup> | **0.08** | **0.31**<br>0.98 | **98.8**<br>85.2 | **93.1**<br>72.6 | **1.88**<br>2.00 | **17.2**<br>26.1 | **97.5**<br>**97.5** | 0.9<br>**0.0** |
|  | Backward<sup>5</sup> | 0.84 | 1.06 | 60.9 | 46.1 | 2.71 | 4.5 | 86.7 | 15.9 |
|  | w/SCWRL<sup>2</sup> | 0.84 | 1.64 | 64.5 | 50.4 | 2.93 | 74.4 | 86.7 | 0.1 |

<sup>1</sup>Side chain $\chi$ -angles were considered accurate when their deviations from experimental values were less than 30 degrees.
<sup>2</sup>Side chains were reconstructed using SCWRL4 after building a backbone structure using other methods (e.g., cg2all, PULCHRA, and REMO).
<sup>3</sup>Chains in multi-chain targets were separately converted to all-atom structures and superposed onto the original Cα-trace.
<sup>4</sup>Conversions of several structures failed because they cannot handle short peptides or were not completed within a reasonable time frame (12 hours). Successful conversions for REMO: 684/720; for ModRefiner: 714/720.
<sup>5</sup>Conversion of one structure failed because short peptides (<3 residues) cannot be handled.

Because all-atom reconstruction via cg2all is achieved with a single forward pass without any iterative optimization, the computational cost is low, on the order of seconds (**Table 3** and *SI Appendix* **Fig. S4**). To better understand the computational performance, some additional analysis is necessary. A reconstruction run with cg2all consists of (1) loading Python libraries, (2) loading a PyTorch model, (3) reading an input PDB file followed by pre-processing, (4) a forward-pass through the model, and (5) writing an output PDB file. Loading the PyTorch model takes around 2.7 seconds, limited by I/O speed. The computational cost for the remaining steps is linearly dependent on the number of residues of the system, with an average of less than 4 seconds for the test set with a single CPU thread and less time when multiple threads or a GPU were used. Using a GPU incurred additional overhead and is not efficient for a single average-size reconstruction, but there is a significant benefit for very large systems or when processing many snapshots with alternate conformations that can be processed simultaneously on a GPU. For a single reconstruction on a single thread, only PULCHRA was faster than cg2all. REMO took about the same total time, whereas other methods required much more time.

**Table 3.**
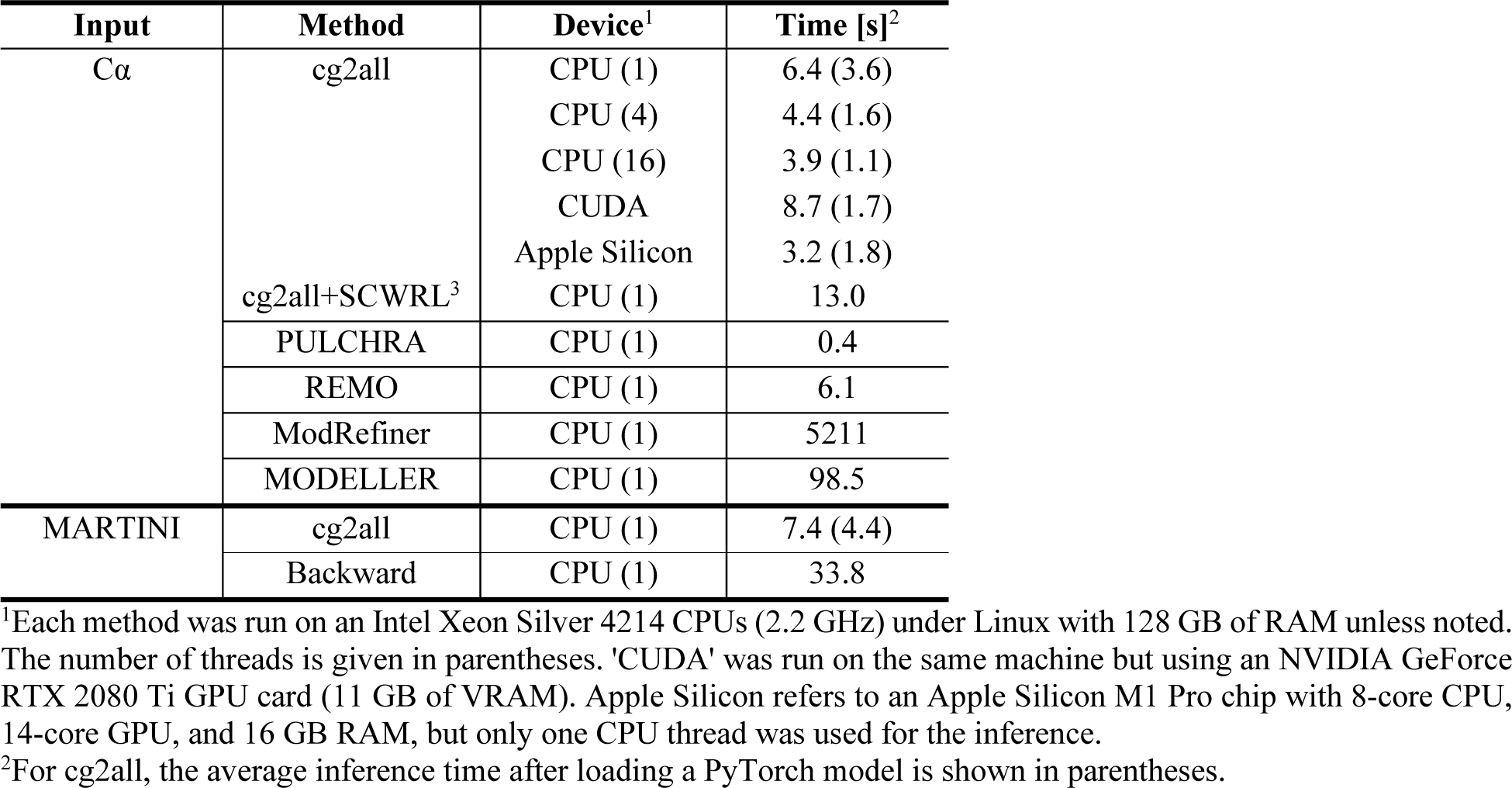
Average timing for all-atom reconstruction with different methods.

### Accurate all-atom reconstruction from simulation models

The reconstruction of all-atom detail from reduced representations of experimental structures, as described in the previous section, may be considered an ideal scenario. We further tested how well all-atom resolution can be recovered from models generated via simulations. Two sets of models were considered. The first set consisted of snapshots that were extracted from all-atom molecular dynamics (MD) simulations; in the second set, the MD-based snapshots were further energy-minimized structures using a residue-based CG model, COCOMO (18). All-atom reconstruction from Cα-traces of MD simulation snapshots was still highly accurate, but the accuracy became slightly worse than for experimental structures, with higher RMSD values of 1.18 Å for heavy atoms and lower side chain torsion accuracies (**Table S1**). Once the snapshots were minimized with COCOMO at the CG level, the all-atom reconstruction RMSD further increased to 1.34 Å (**Table S1**) with respect to the initial all-atom MD snapshots. However, that may be expected because the CG minimization by itself resulted in deviations of 0.30 Å for the Cα positions from the initial all-atom MD snapshots. In either case, the reconstructed all-atom models had again low clash scores and very low rotamer outliers and were much closer to the reference atomistic structures than those generated with other methods (*SI Appendix* **Table S1**). Thermal fluctuations in the simulations led to broadened bond geometry distributions (*SI Appendix* **Fig. S5**) and more rotamer outliers compared to experimental structures. However, since cg2all was trained to generate experimental structure-like conformations, the broader distributions were not completely reproduced (*SI Appendix* **Fig. S5**). This explains at least in part the slightly lower reconstruction accuracy for simulation-based models.

A related practical question is whether the reconstructed models from cg2all are more suitable for starting atomistic simulations. The reconstructed all-atom structure was suitable for further usages such as all-atom MD simulations. Larger systems require extensive computational cost to get their equilibrated system or systems in desired states via all-atom MD simulations. Alternatively, one may attempt to reach an equilibrium state such as LLPS formation described in **Fig. 2C** using CG simulations (18) and continue atomistic simulations from the state. We briefly examined the approach by performing atomistic MD simulations starting from reconstructed all-atom structures (*SI Appendix* **Fig. S3**). We carried out an atomistic MD simulation for 50 ns and minimized the conformation from the last snapshot using the CG model COCOMO as a hypothetical CG simulation result for which the atomistic MD snapshot serves as a reference. The COCOMO minimized structure was converted to an all-atom structure using our method or PULCHRA (28), respectively. Then, the converted structures were equilibrated again and continued atomistic simulations. The simulation results were compared with another set of simulations that simply continued simulations from the last snapshot. We hypothesized that the protein structure would quickly show instabilities at the beginning of the simulation if the conversion was not producing models of sufficient quality. After conversion with cg2all, the protein structure remained stable and well-folded during the first 10 ns, as the continued simulation did. The average Cα-RMSD to the initial conformation was 1.98 Å after 10 ns (*cf.*, 1.35 Å for the simulations from the last snapshot). On the other hand, with the reconstructed structure by PULCHRA, due to steric clashes, conformations deviated from the initial conformation significantly, starting at early stages of the simulations (2.94 Å Cα-RMSD with respect to the initial conformation on average after 10 ns). Consequently, residue-wise fluctuations (RMSF) throughout the atomistic simulation were very similar between the sets of simulations from the last snapshot and the model by our method, while an initial model from PULCHRA resulted in significantly higher fluctuations due to initial destabilization caused by steric clashes.

### Adding atomistic detail to low-resolution models

The analysis so far shows that all-atom details can be captured essentially within experimental uncertainties at much reduced representations, up to a single bead at the center of mass of an amino acid. This is possible by drawing on the vast knowledge about protein structures via state-of-the-art machine learning. In turn, this means that all-atom detail can be provided with high confidence for models that are initially only available at a CG level. Examples where cg2all may be used in practice are shown in **Fig. 2** and discussed more in the following.

Low-resolution cryo-EM structures are often reported only at the Cα level. All-atom detail can be reconstructed quickly via cg2all (**Fig. 2D**). For a cryo-EM experimental structure (PDB ID: 3iyg (39)) with a resolution of 4.0 Å the all-atom structure with 4,134 residues or 64,192 atoms was generated within 11.9 seconds, fast enough to be done on-demand when working with such structures. The generated structure had a higher density map correlation of 0.723 than that of the original Cα-trace, 0.647, but there were some clashes in the atomistic model, presumably because Cα atoms for some residues were packed too tightly in the original structure. Therefore, it may be possible to use the all-atom reconstruction as an indicator of issues with the low-resolution model itself.

Residue-level CG models, such as COCOMO, are increasingly being used to simulate very large systems over long time scales, for example to study protein-protein interactions or liquid-liquid phase separation. Again, cg2all can provide atomistic detail from the CG models (**Figs 2C** and **2E**). Using this approach, we could obtain an all-atom structure from a snapshot of a CG model of a condensate formed by IDPs. In the example, a 14,112-residue CG system was converted to an all-atom structure with 183,624 atoms in just 33.5 seconds. This rendered detailed atomic interactions between peptides (*e.g.*, salt bridges between charged side chains) inside the condensate. We note that a fully atomistic simulation of the condensation process is so far impossible to carry out.

Machine-learning based conformational ensemble generators such as FoldingDiff (41) or idpGAN (14) may be limited to output consisting of Cα traces due to resource constraints, but all-atom models can be obtained rapidly via post-processing by cg2all (**Fig. 2F**). Using a GPU with a large batch size allowed us to convert 25,000 IDP conformations generated via idpGAN to all-atom detail in just 64.6 seconds. Consequently, a practical strategy for the rapid generation of conformational ensemble via machine-learning may be to focus on generating ensembles only at the CG level and leave it up to an all-atom reconstruction scheme as presented here to obtain atomistic ensembles.

Finally, since cg2all can efficiently parallelize all-atom reconstructions on GPUs, entire CG MD simulation trajectories could be rapidly converted to atomistic detail. For example, the conversion of 10,000 frames of a 1,824-residue system takes 1,774 seconds on a GPU with a batch size of 4 (*SI Appendix* **Video S1**). This allows not just the consideration of all-atom detail when sampling with CG models, but also, *vice versa*, suggests that CG models could be used for lossy data compression. This has been proposed before based on the higher-resolution CG model PRIMO (42), but much greater compression can be achieved if only a single particle per residue is used. For example, a 3.4 GB all-atom trajectory in the DCD format could be compressed into a 210 MB trajectory with single beads, such as Cα or center-of-mass, resulting in a 94% compression ratio. Such high degree of compression could greatly facilitate the public sharing of extensive atomistic trajectories that otherwise remains a significant resource challenge (43, 44).

### Multi-scale sampling for rapid cryo-EM refinement

To further demonstrate the potential of cg2all, we turn to the refinement of models against cryo-EM densities. A typical challenge involves the flexible fitting of initial models from crystallography or structure prediction to intermediate- to low-resolution density maps, for which direct atomistic model building is difficult due to insufficient information (45). The most effective methods to date employ sampling via atomistic simulations, such as the MDFF protocol (46). This approach is successful but may take on the order of hours to days because of the computational cost of the simulations. Here we explore the sampling of CG models guided by a density map correlation energy function based on reconstructed all-atom representations that is possible with cg2all. Sampling at the CG level avoids the kinetic barriers that hinder sampling at the atomistic level, whereas using an energy penalty based on atomistic reconstructions ensures that the optimized CG model is maximally compatible with the experimental data.

The multi-scale approach based on cg2all outperformed local optimization protocols such as energy minimization at the all-atom representation using an atomistic energy function or energy minimization at the CG level using a CG energy function alone across the entire range of map resolutions (**Figure 3 and** *SI Appendix* **Figs. S13 and S14**). With a high-resolution (3 Å) electron density map, local energy minimization of an all-atom model is trapped in a local energy minimum because of a rugged energy landscape. On the other hand, local energy minimization of a CG model cannot exploit the high-resolution information from the electron density map. However, our multi-scale approach effectively optimizes structures by minimizing at the CG level where kinetic barriers are low or absent while still targeting the high-resolution data via all-atom reconstruction. The optimized structures obtained via the cg2all-based multi-scale approach reached comparable C⍺-RMSD values to the full MDFF protocol (0.36 vs. 0.35 Å), slightly lower cross-correlation coefficients (CCC, 0.866 vs. 0.881) and slightly larger heavy atom-RMSD values (0.88 vs. 0.74 Å). Importantly, the similar accuracy with cg2all vs. MDFF is achieved in much shorter time (minutes vs. hours). An example for the high model accuracy that can be achieved with cg2all is shown in **Fig. 4**. In the example, several regions in the initial AlphaFold2 model located outside of the 5 Å resolution electron density (indicated by red arrows) with a heavy-atom RMSD of 2.26 Å and a CCC of 0.829. Using the MDFF protocol with the electron density map, the model was optimized to a heavy-atom RMSD of 0.80 Å and has a higher CCC of 0.941. Our multi-scale approach optimized the model to a comparable accuracy, a heavy-atom RMSD of 0.85 Å and a CCC of 0.936, even though it performed the actual optimization at the CG-level. We note that our multi-scale approach achieved the comparable accuracy in 8.9 minutes, while the MDFF protocol took 8.8 hours.

**Figure 3.**
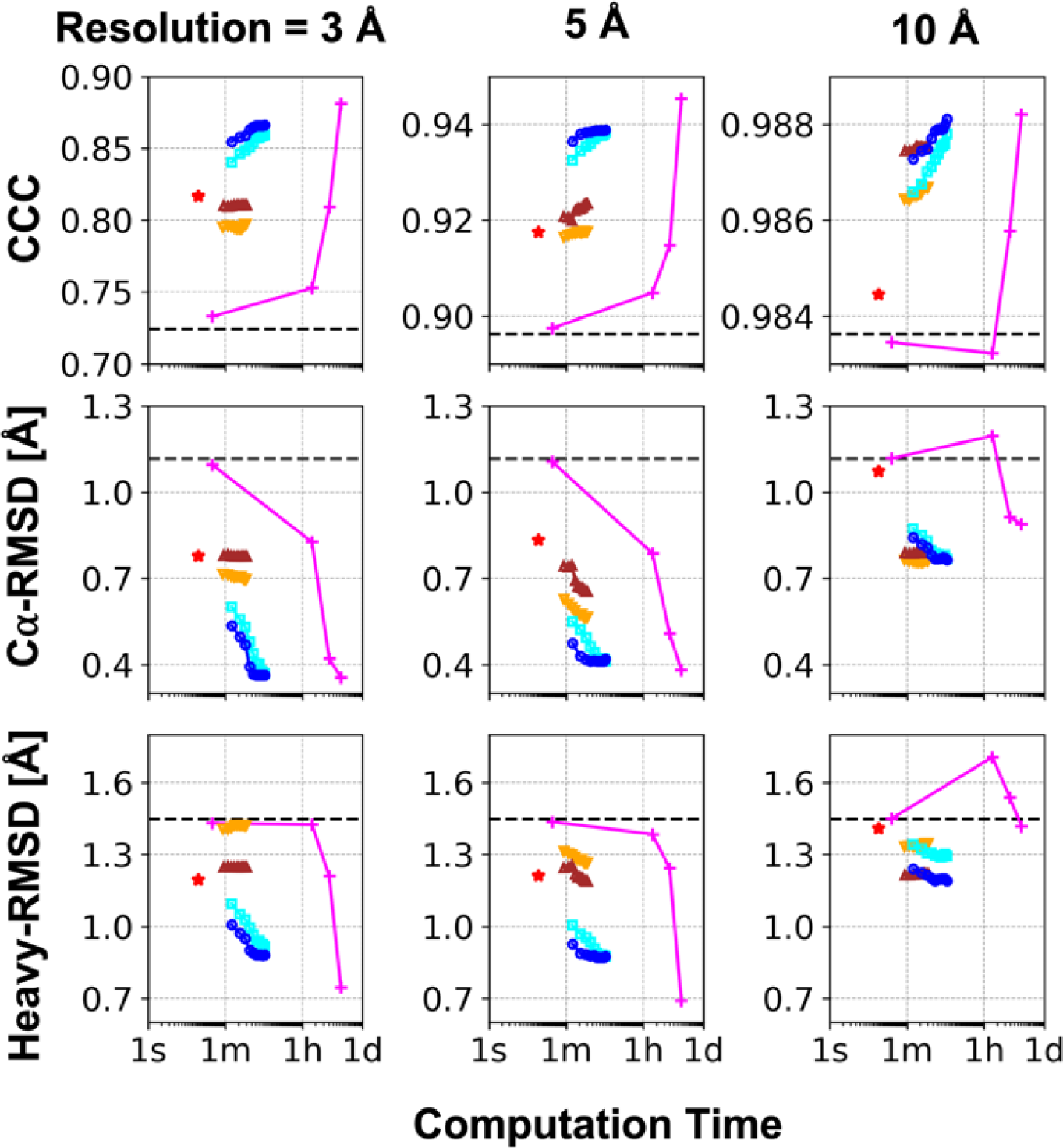
Refinement of AlphaFold models against cryo-EM density maps. Model quality in terms of cross correlation coefficient (CCC) against target cryo-EM density maps, Cα and heavy atom RMSDs were analyzed as a function of computation time for several protocols: (1) optimizations using cg2all models for residue center-of-mass model (blue circles) and Cα-trace (cyan circles) followed by all-atom energy minimization, (2) optimization at the residue center-of-mass (brown triangles) or Cα-trace representation (orange triangles) followed by all-atom energy minimization, (3) all-atom energy minimization only (red star), and (4) MDFF samplings (magenta ‘+’). The initial model qualities are shown as black dashed lines.

**Figure 4.**
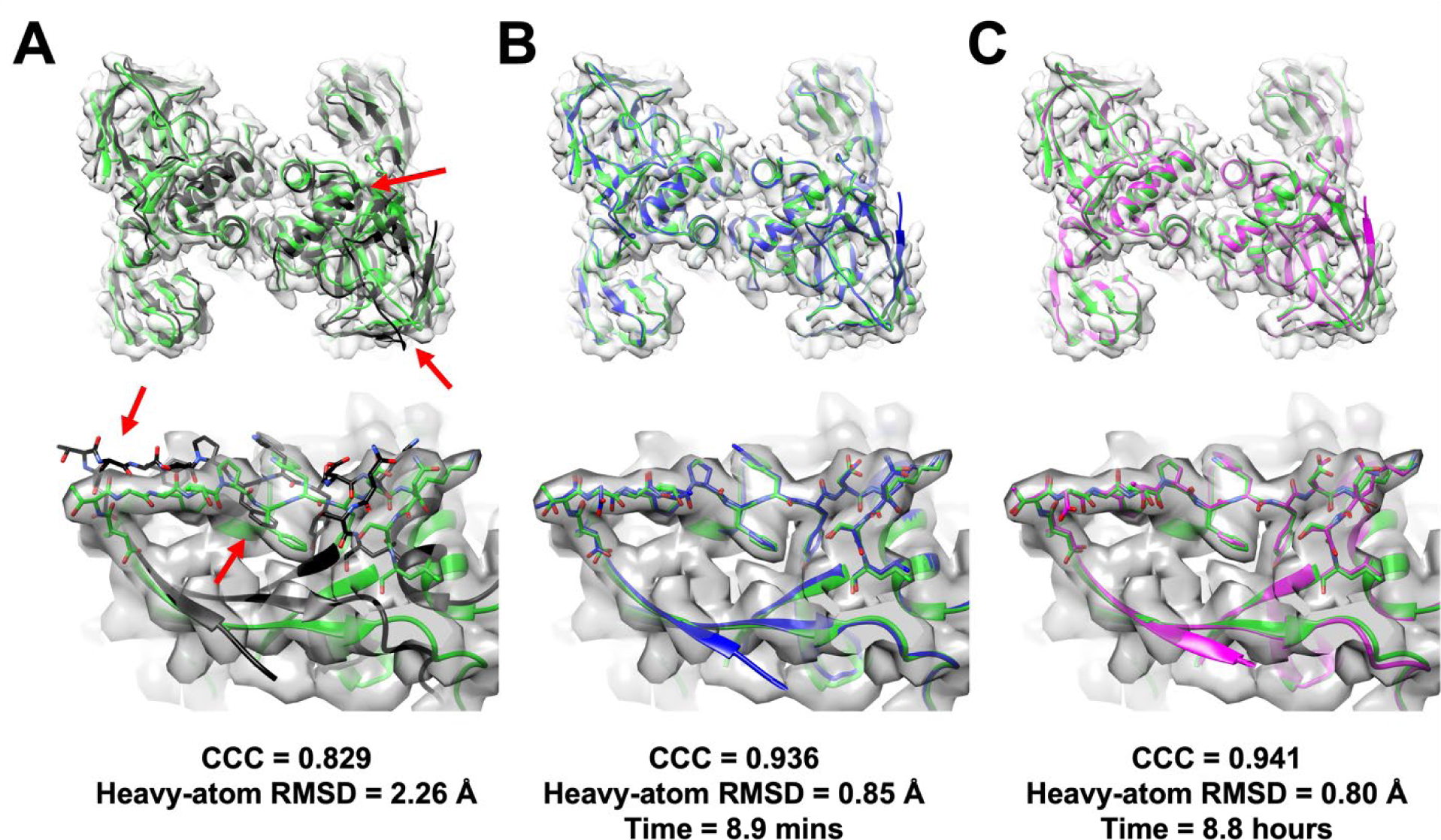
Refinement against a cryo-EM density map for 3isr. The target tetrameric structure (PDB ID: 3isr (49)) is shown in green cartoon representation. Its synthetic electron density map was created at 5 Å resolution using “molmap” command in UCSC Chimera (50), which is based on EMAN2’s pdb2mrc program (51), and depicted as transparent grey surface at a density level of 0.3. The initial model structure generated by AlphaFold-Multimer (52) (A), its optimized structure using cg2all model for Cα-trace and followed by all-atom minimization (B), and another optimized model via all-atom minimization using MDFF (C) are shown in black, blue, and red. Overall tetrameric structures and a region where significant deviations (indicated by a red arrow) were optimized are shown in the top and bottom panels.

When the electron density map has much lower resolution, such as 10 Å, optimization at the all-atom level becomes less effective, even using MDFF, whereas the CG-based optimized refinement, via cg2all, still allows structure refinement, and still within minutes. This opens up the possibility for high-throughput model refinement of many lower-resolution maps, for example to fit models to maps of dynamics conformational ensembles captured via cryo-EM.

In this proof-of-concept demonstration, we employed a naïve CG energy function that only prevents severe clashes between CG beads. Distance restraints between CG beads were applied to keep the protein structure folded, however, this also limited the potential for improvement as it prohibited partial structure unfolding and refolding (47, 48). In future work, this will be addressed by introducing a more sophisticated CG energy function that is capable of not only maintaining folded structures but also allowing significant structural changes.

## CONCLUSIONS

The results presented here show that all-atom details of proteins can be captured essentially within experimental uncertainties with only a single bead per amino acid, especially when placed at the residue center of mass, but perhaps also with a more traditional Cα-trace representation. This is possible now because of advances in machine learning that allow vast information from known structures in the PDB, to be applied towards different objectives, in this case, the accurate reconstruction of all-atom features from low-resolution models.

The approach taken here was initially motivated by recent advances in protein structure prediction methods, but it is different in terms of input as well as the final objective. Sequence alignments or template libraries are not used here, instead a lower-resolution model serves as input. On the other hand, even although the ultimate goal of providing physically realistic, accurate atomistic structures is essentially the same, structure prediction methods aim at providing the best model for the likely native state whereas the method introduced here aims at generating atomistic detail for any conformation, whether energetically favorable or not. This suggests that recent advances in machine learning have broad implications for structural biology that reach far beyond just the prediction of native structures from sequence.

There are immediate applications in adding accurate atomistic detail to CG representations, within the limitation that cg2all learned to reproduce the most likely time- and ensemble-averaged conformations for otherwise dynamic residues. Low-resolution protein structure models based on experiments as well as CG models from simulations can be interpreted in atomistic detail. This is especially relevant for the generation of structures and ensembles via machine learning where the addition of atomistic detail often presents a significant burden during model training.

Finally, an important feature is that deterministic neural network architectures are not just very efficient but allow gradients to be back-propagated all the way from the final output (*i.e.* atomistic conformations) to the input (*i.e.* CG conformations). In essence, this provides an avenue for tightly coupled bidirectional multi-scaling. While there are many possible directions that we expect could be pursued in the future, we highlight one possible application: refinement of models against cryo-EM densities based on sampling at a CG level but with energy penalty functions evaluated at the atomistic level from reconstructed all-atom conformations. Just for this application we demonstrate comparable performance to much more time-consuming atomistic simulations with computational costs that are an order of magnitude less. We expect that many other applications can be realized in a similar manner.

## METHODS

### Datasets

Two sets of experimental structures were used for training, validation, and test. The first one, PDB 6k, originated from the Top8000, which is a curated protein structure dataset only with high-resolution X-ray structures (53). The second set, PDB 29k, was curated by culling X-ray structures with high-resolution (≤ 3.0 Å) and lower R-value (≤ 0.3) using PISCES on September 9, 2022 (54). To test the model’s applicability to non-experimental an ensemble of structures was generated by performing all-atom MD simulations. In addition to the all-atom MD simulation snapshots, their energy-minimized conformations using the COCOMO Cα-trace CG model were introduced to mimic sampling at the CG level.

### Model architecture

The neural network models for the conversion of a CG model to an all-atom structure were based on an SE(3)-equivariant architecture (**Fig. 1**), with three modules: an initialization module to encode input features, an interaction module based on an SE(3)-transformer (55), and a structure module to generate the final output. A protein structure was described as a homogeneous bi-directional graph. Each protein residue corresponds to a graph node. For a residue-level CG model (one bead per residue), all beads were used as nodes. For a MARTINI model, BB beads were considered as representative beads for residues. Nodes were connected by edges if the distance between the representative beads of a residue was shorter than a distance cutoff (*e.g.*, 10 Å). As input node features, scalar (*l*=0) and vector (*l*=1) properties extracted from geometric relationship among neighboring nodes were used (*SI Appendix* **Fig. S7**). More details are given in the *SI Appendix*.

## DATA AVAILABILITY

The source code and model parameter files of the method are available at https://github.com/huhlim/cg2all. It can be locally installed using a PIP command, “pip install git+http://github.com/huhlim/cg2all”. Alternatively, demo services are available at https://huggingface.co/spaces/huhlim/cg2all and https://colab.research.google.com/github/huhlim/cg2all/blob/main/cg2all.ipynb. A Google Colab notebook for local optimization with cryo-EM density map is available at https://colab.research.google.com/github/huhlim/cg2all/blob/main/cryo_em_minimizer.ipynb.

## ACKNOWLEDGEMENTS

Funding was provided by the National Institute of Health (NIGMS) grant R35 GM126948.

## AUTHOR CONTRIBUTIONS

LH and MF designed the research, LH performed and analyzed the work, and LH and MF jointly wrote the manuscript.

