## Supplemental Text, Figures, and Tables for "One particle per residue is sufficient to describe all-atom protein structures"

**This PDF file includes:**

Supplementary methods
Algorithm S1
Figures S1 to S14
Table S1
Video S1 Caption
References for SI reference citations

#### Supplementary Methods

##### *Experimental dataset details*

Two sets of experimental structures were clustered with a maximum mutual sequence identity of 70%, and one structure for each cluster was left. The original Top8000 (1) and the PISCES (2) sets consisted of a single chain per Protein Data Bank (PDB) entry, however, we used all protein chains in each PDB entry instead. PDB entries with more than 1,200 residues were excluded. As a result, there were 7,130 and 30,354 entries in PDB 6k and 29k sets, respectively. To have common subsets for model validation and test, 720 entries were randomly selected among the common entries for each validation and test sets. The remaining 5,690 and 28,914 structures were used as PDB 6k and 29k training sets, respectively.

The Top8000 structures were further analyzed to obtain statistics of bonded geometries for building secondary structure-dependent rigid body (SS-dep) blocks. Three-state secondary structure for every residue was assigned by the DSSP algorithm (3) using the MDTraj Python library package (4). Bond lengths, angles, and improper dihedral angles defined by amino acid topologies of the CHARMM36m force field (5) were calculated. Their averaged values for each secondary structure type were then used to build SS-dep blocks.

##### *Simulation dataset details*

We sampled an ensemble of structures by performing all-atom molecular dynamics (MD) simulations using OpenMM (6). A protein structure was placed at the center of a periodic rectangular box with at least 10 Å distance from any protein atom to any dimension of the box edges. The remaining space in the simulation box was filled with the CHARMM version of TIP3P water molecules (7). Some water molecules were randomly replaced with sodium or chloride ions to neutralize the system and achieve a total ion concentration of 0.015 M. The CHARMM36m force field (5) was applied to describe the system throughout the series of simulations. The system was locally minimized with the l-BFGS-b algorithm (8) in the presence of harmonic positional restraints on every Cα atom with a force constant of 0.5 kcal/mol/Å$^2$. Then, the systems were gradually heated to 298.15 K and equilibrated via Langevin dynamics simulations for 1 ns with a friction coefficient of 0.01/ps and a 2-fs integration time step. The NVT ensemble and the NpT ensemble at 1 bar with a Monte Carlo barostat were applied during the heating and equilibration steps, respectively. An ensemble of protein conformations was sampled from a 50 ns-long Langevin dynamics simulation with a friction coefficient of 1/ps and a 2-fs integration time step at 298.15 K and 1 bar. Five snapshots of protein conformations were picked up for further tests by selecting frames for every 10 ns. In total, 3,600 all-atom conformations were generated for 720 test set structures for further tests.

During minimization with COCOMO (9), harmonic positional restraints were applied to every bead of the CG model with a force constant of 1.0 kcal/mol/Å$^2$ to keep the original all-atom conformations as the respective ground truth conformations. Upon minimization, the conformations were distorted from their all-atom structures by 0.30 Å on average.

##### *Input feature details*

In total, 57 (17 from the local geometries and 40 from the residue type embedding) scalar and four (or more for multiple site CG models) vector node features and three scalar edge features were used as input features. Pseudo-bond angles and -torsion angles were encoded using cosine and sine functions to account for periodicity. The number of neighboring nodes (*e.g.*, < 10 Å) and the presence of a previous or next residue (to distinguish terminal residues) were added as scalar features. The residue type was converted to 40 scalars via a trainable embedding layer and concatenated with the scalar features. For vector features, unit bond vectors as described in *SI Appendix* **Fig. S7** were used For conversion from multiple site coarse-grained models such as -a MARTINI model (10), vectors from a BB bead to SC beads were additionally used to incorporate side chain information. The edge connection type (connection via a peptide bond, an inter-residue contact through space, or a disulfide bond) were used via one-hot encoding as edge features.

***Neural network model details***

At the core of SE(3)-equivalent neural network model, SE(3)-Transformers architecture (11) was adopted. The input node features were processed via N (=4 for the baseline model) linear layer blocks to produce 64 scalar and 32 vector values. A LayerNorm (12) and an exponential linear unit (ELU) activation function (13) were used prior to a linear layer in a block. Until this step, interactions between nodes were not considered yet. In the interaction module, M (=4 for the baseline model) SE(3)-Transformers blocks with a LayerNorm and an ELU activation function were facilitated to communicate between nodes. For the SE(3)-Transformers, we used eight attention heads and 32 hidden channels of scalars ($l$=0), vectors ($l$=1), and rank-two tensors ($l$=2).

We used four linear layers and four SE(3)-Transformers blocks. Alternatively, fewer and greater numbers of blocks were examined to evaluate performance dependencies in the model size. We adopted ELU activation functions as replacements of rectified linear unit (ReLU) functions (14), which were used for the original SE(3)-Transformers work (11). For hidden features of the SE(3)-Transformers blocks, features up to the degree of 2 were passed within a block, and a lower value (1) was also tested. For a protein structure, we used a subgraph with a crop size of 256 residues as the baseline and tested if a larger crop (384 residues) could improve the performance. Secondary structure-dependent rigid-body blocks were not used for the baseline, but they were adopted as optional features.

***Structure module details***

The structure module further processed the output of the interaction module to predict values for building all-atom structures via N (=4, for the baseline model) linear layer blocks. An all-atom structure was built using the predicted values. For this process, a similar procedure that is used by AlphaFold2 was adopted (15). AlphaFold2 used rigid-body blocks of backbone atoms (N, Cα, and C atoms), the backbone oxygen atom, and side chain heavy atoms that were segmented by rotatable torsion angles. During the step of structure building, backbone rigid-body blocks for residues were oriented first using predicted translations (***t***) and three-dimensional rotations (**R**). The remaining blocks were placed in the order of the bond connectivity from the backbone block to the tip of each side chain by rotating them by predicted torsion angles ($\phi$, $\psi$, and $\chi$s). In our work, we extended the procedure to build all-atoms including hydrogen atoms. In addition, secondary structure-dependent rigid body blocks were used as we found that some blocks have distinguished bonded geometries (bond lengths and angles and improper dihedrals) depending on the secondary structure. Thus, secondary structure prediction (SS) was also made by the module. We note that we used a 6D representation (16), which describes three-dimensional rotation using two unit vectors and the Gram-Schmidt process, instead of the quaternion-based approach used by AlphaFold2, as it gave better performance in terms of accuracy and convergence during training for many SO(3) prediction tasks because of continuity in the rotation representations (16). For the final step, both N- and C-termini were patched by replacing backbone hydrogen (HN) and oxygen (O) with the terminal amino ($NH_3$-) and carboxyl (-$COO^-$) groups, respectively, using the predicted position of the atoms.

##### *Model training*

Each model was trained for 300 and 120 epochs with the PDB 6k and 29k training sets, respectively, using a batch size of 4. This corresponds to 426,600 and 867,360 total training steps. An input data graph with more than 256 nodes for training was randomly cropped to a consecutive 256 residue graph by obtaining its subgraph. An Adam optimizer was used with learning rates of 0.01 for parameters for scalar features of the structure module and 0.001 for the others. The learning rates were linearly increased for the first ten epochs and then exponentially decreased with a multiplicative factor of 0.995. Clipping was used to prevent extreme gradients, and gradient checkpointing reduced memory consumption. Models were implemented in PyTorch, and they were trained using PyTorch Lightning (17). The training of models was carried out on two NVIDIA GeForce RTX 2080 Ti GPU cards (11 GB of VRAM) with Distributed Data Parallel (DDP) to use multiple GPUs and eight CPU threads. We trained three models for each model variant to obtain statistics.

The progress in learning protein structural features by the Cα-trace model proceeded in the order of distance from the Cα atom (**Fig. S8**). When progress in the recovery of structural features was tracked for every 10 epochs, we observed that backbone-related features such as the Ramachandran angle, the result of translation and three-dimensional rotation of backbone rigid-body blocks, were saturated during the earlier epochs of the training. At epoch 10, the Ramachandran map already resembled that from experimental structures, and it changed little afterwards. On the other hand, learning side chain torsion angles required many more epochs. At epoch 20, predictions of $\chi_1$ angles became reasonably accurate, and the model started to learn $\chi_2$ angles and a little bit of $\chi_3$ angles. At epoch 60, more states of $\chi_2$ angles were captured, and there was progress in $\chi_3$ angle predictions. At the end of the training at epoch 120, learning of most structural features converged. Many structural features were learnt by the model, however, a few torsion angles such as those for Arg/Lys $\chi_4$ angles could not be learnt in the end. Consequently, the loss in structural information upon the conversion from all-atom structure to Cα-trace was most significant for structural features that were far from the Cα atom. Consequently, backbone features could be learnt quickly, while torsion angles farther from the Cα atoms were slow and sometimes incomplete. In contrast, the learning progress with the residue-center-of-mass model, which contains richer input information, was much faster overall and more complete than that with the Cα-trace model. (**Fig. S9**) Most structural features started to converge at epoch 20 and were almost completed at epoch 60.

##### *Loss function*

Models were trained with a loss function (**Eq. 1**) that consisted of a training data-dependent loss and a data-independent (physics-based) loss:

$$L = \begin{matrix} 5.0\,L_{FAPE,C\alpha} + 1.0\,L_{BB} + 1.0\,L_R + 1.0\,L_{backbone\ geometry}\ (+0.1\,L_{SS}) & (Data-dependent) \\ +5.0\,L_{torsion} + 1.0\,L_{v_{cntr}} & \\ +5.0\,L_{atomic\ clash} + 0.1\,L_{torsion\ energy} + 1.0\,L_{side\ chain\ geometry} & (Physics-based) \end{matrix} \quad (1)$$

The FAPE, Cα loss ($L_{FAPE,C\alpha}$) was first introduced by AlphaFold2 (15), and we used a variant that uses only Cα atoms for its evaluation with a distance clamp value of 10 Å. The backbone loss ($L_{BB}$) also contributed to correctly placing backbone rigid bodies (**Eq. 2**).

$$L_{BB}(\,r_{BB}^{truth}) = \left|r_{BB} - r_{BB}^{truth}\right| + \left|1 - \overrightarrow{u_{C\alpha\to N}} \cdot \overrightarrow{u_{C\alpha\to N}^{truth}}\right| + \left|1 - \overrightarrow{u_{C\alpha\to C}} \cdot \overrightarrow{u_{C\alpha\to C}^{truth}}\right| \quad (2)$$

where $\vec{u}$ represents a unit vector. The loss function for three-dimensional rotation representation consisted of the similarity of two vectors for the 6D representation (16) against their truths and an auxiliary term that ensures their vector sizes close to 1. (**Eq. 3**)

$$L_R\left(\overrightarrow{R_0}, \overrightarrow{R_1} \,|\, \overrightarrow{R_0^{truth}}, \overrightarrow{R_1^{truth}}\right) = \left|\left(\overrightarrow{R_0} - \overrightarrow{R_0^{truth}}\right) + \left(\overrightarrow{R_1} - \overrightarrow{R_1^{truth}}\right)\right| + 0.01\left(\left|\left\|\overrightarrow{R_0}\right\| - 1\right| + \left|\left\|\overrightarrow{R_1}\right\| - 1\right|\right) \quad (3)$$

where $\overrightarrow{R_0}$ and $\overrightarrow{R_1}$ are vectors for the 6D representation. A loss function helped the model having correct bonded geometries for backbone atoms (**Eq. 4**). It penalized deviations of peptide bond distances ($b_{C,+N}$) and peptide bond including bond angles ($\theta_{C\alpha,C,+N}$ and $\theta_{C,+N,+C\alpha}$).

$$L_{backbone\ geometry}\left(b_{C,+N}, \theta_{C\alpha,C,+N}, \theta_{C,+N,+C\alpha} \,\middle|\, b_{C,+N}^{truth}, \theta_{C\alpha,C,+N}^{truth}, \theta_{C,+N,+C\alpha}^{truth}\right) = \left|b_{C,+N} - b_{C,+N}^{truth}\right| + \frac{1}{2}\left(\left|\theta_{C\alpha,C,+N} - \theta_{C\alpha,C,+N}^{truth}\right| + \left|\theta_{C,+N,+C\alpha} - \theta_{C,+N,+C\alpha}^{truth}\right|\right) \quad (4)$$

When the secondary structure-dependent rigid body blocks were used, a cross entropy loss for secondary structure prediction was additionally used.

Moreover, there were two loss functions for correctly reconstructing side chain atoms. The torsion angle loss tried to reduce discrepancies between the predicted torsion angles ($\{\theta_k\}$) and their truth values ($\{\theta_k^{truth}\}$) (**Eq. 5**).

$$L_{torsion}\left(\{\theta_k\} \,|\, \{\theta_k^{truth}, \theta_k^{truth,alts}\}\right) = \sum_{\theta_k\ is\ defined}\left(1 - max\left(cos\left(\theta_k - \theta_k^{truth}\right), cos\left(\theta_k - \theta_k^{truth,alts}\right)\right)\right) \quad (5)$$

Some side chain torsion angles that have periodicity: $\chi_2$ angles of Phe and Tyr, methyl groups in Ala, Ile, Leu, Met, Thr, and Val, the side chain amino group in Lys, and the side chain carboxyl groups in Asp and Glu. For those torsion angles, alternative truth values ($\{\theta_k^{truth,alts}\}$) due to their periodicity were considered for the calculation. Furthermore, solvent-exposed side chains can have multiple valid conformations, while their experimental structures usually presented only one of them. For the training of models that convert from a Cα-based model, putative side chain conformations were generated prior to the training and used as additional $\{\theta_k^{truth,alts}\}$. The procedure for putative side chain conformation generation is described in more detail below. The other loss function for side chain atoms depended on vectors from the Cα atom to the center of mass of a residue ($\overrightarrow{v_{cntr}}$). Their deviation in their vector size to their truth values and their cosine similarity was used to learn side chain orientations at a lower resolution.

$$L_{v_{cntr}}\left(\overrightarrow{v_{cntr}^{truth}}\right) = \left|\left\|\overrightarrow{v_{cntr}}\right\| - \left\|\overrightarrow{v_{cntr}^{truth}}\right\|\right| + \left|1 - \overrightarrow{u_{cntr}} \cdot \overrightarrow{u_{cntr}^{truth}}\right| \quad (6)$$

In addition to these data-dependent loss functions, three physics-based terms were introduced to improve the geometric properties of the reconstructed models. These terms relied on the CHARMM36m force field (5). As two atoms rarely overlapped within the sum of their atomic radii, atomic clashes were penalized (**Eq. 7**).

$$L_{atomic\ clash}(r) = \sum_{i<j, r_{ij}<14Å} \sum_{a\in i} \sum_{b\in j\ exclude\ 1-2,3,4\ pairs} \sqrt{\varepsilon_a \varepsilon_b} \times \left(\min(0, d_{ab} - \sigma_a - \sigma_b)\right)^2 \quad (7)$$

where $\varepsilon$ and $\sigma$ are the depth of the potential and the distance at which the potential becomes zero in the Lennard-Jones potential. Torsion energy terms ($L_{torsion\ energy}$) of the force field was applied to penalize disfavored torsion angles (**Eq. 8**). Torsion angles that can be defined within a residue were considered for evaluation, thus, torsion angles that span two consecutive residues (*e.g.*, ψ, ϕ, and ω angles) were not subjected to the loss functions. In order to preserve torsion angle distributions, a torsion energy clamp ($E^{clamp}_{torsion\ energy}$) was used with a value of 0.6 kcal/mol.

$$L_{torsion\ energy}\ (\{\theta_k\}) = \sum_{\theta_k\ is\ defined} \max\left(0, E_{torsion\ energy}(\theta_k) - E^{min}_{torsion\ energy}(\theta_k) - E^{clamp}_{torsion\ energy}\right) \quad (8)$$

The final physical loss term was applied for two types of bonds that connect rigid body blocks in special ways, namely for proline ring closure and disulfide bonds. For these bonds, equilibrium bond lengths of 1.455 and 2.029 Å, respectively, were targeted (**Eq. 9**).

$$L_{side\ chain\ geometry}\left(b_{Pro,N,CD}, b_{SSBOND}\right) = \left|b_{Pro,N,CD} - b^{o}_{Pro,N,CD}\right| + \left|b_{SSBOND} - b^{o}_{SSBOND}\right| \quad (9)$$

***Augmentation of putative side chain conformations***

Alternative possible side chain conformations were generated prior to the training using first SCWRL4 (18), followed by REDUCE (19) and local energy minimization using the CHARMM36m force field (5). Experimental structures, especially those determined by X-ray crystallography, have only one conformation in PDB in most of the entries even though there can be alternative coordinates. However, as proteins are not static molecules, they can have diverse conformation especially for solvent-exposed side chains. As such, reconstruction to those alternative conformations should not be penalized unless they are unfavorable. Because we aimed to reconstruct an all-atom conformation including side chains for a given Cα-trace of a protein, we generated putative side chain conformation. For a protein structure, side chain structures were predicted on the protein's backbone using SCWRL4, which uses a rotamer library and optimizes combinations of rotamer states. Then, all hydrogens were attached to the predicted structure with an optimization of torsion angles for the side chain amide groups in Asn and Gln and the imidazole ring in His. Finally, the structures were subjected to energy minimization using the l-BFGS-b algorithm for up to 1,000 steps with the CHARMM36m force field using OpenMM (6). To prevent extensive deviation of backbone positions, harmonic positional restraints were applied on every N, Cα, C, O, and Cβ atoms with a force constant of 1.0 kcal/mol/Å$^2$. Torsion angles from the energy minimized structure were used as an additional set of $\left\{\theta_k^{truth,alts}\right\}$ for the torsion angle loss (**Eq.** 5).

***Hyperparameter optimization and ablation studies***

Several features were introduced in our neural network models and their training, and their contributions were evaluated via ablation studies. (SI Appendix **Figs S10** and **S11**) First of all, we used a hybrid loss function that consisted of data-dependent and -independent (or physics-based) loss functions (**Eq. 1**). The physics-based loss functions were introduced to learn the characteristics of a protein molecule more efficiently and to complement insufficient data points. The atomic clash loss penalized inter-atomic clashes and effectively lowered the clash score. Side chain modeling as rotamer outliers could be suppressed by introducing the torsion energy loss function (**Fig. S12**). Without the torsion energy loss function, side chain rotamer states could not be clearly separated, and some inferences resulted in rotamer outliers. Average MolProbity scores (20) by models with and without those physics-based loss functions were different by

0.254 (2.292 vs. 2.546). Furthermore, side chain conformations were augmented prior to the model training to account for their putative heterogenic conformations due to their conformational flexibility. Because some side chains (especially solvent-exposed ones) can have alternative conformations in addition to the experimentally resolved one, both conformations should not be penalized unless there are atomic clashes. This side chain augmentation additionally helped suppress generating rotamer outliers (**Figure S12**). If there were side chains with very similar input features but in different rotamer states in the training set, models could be trained to predict in the average of the rotamer states unless the side chain augmentation was used, and this could result in the inference of rotamer outliers. For example, for a homodimer with pseudo-$C_2$ symmetry, some side chain conformations may vary in different monomers, while overall Cα-traces were almost identical. As illustrated in **Fig. S12B**, if an Arginine from a monomer has a $\chi_1$ angle of -180 degree, while the other Arginine in the symmetry has a $\chi_1$ angle of -60 degree, training with these data would result in predicting their averaged value, -120 degree, which is a rotamer outlier. It is because the averaged torsion angle loss has a minimum at the value. With the consideration of both possible conformations in the torsion angle loss function via the side chain augmentation, the loss function has multiple minima (*e.g.*, -180 and -60 degrees in this example), and the inference of rotamer outliers is diminished.

We tested two neural network parameters that were related to the amount of information passed between layers: 1) the choice of the activation function (ELU vs. ReLU); and 2) the maximum degree for the SE(3)-Transformers (*l*=2 vs. 1). When the ELU activation function was replaced with the ReLU function, which was originally used in SE(3)-Transformers, the overall quality of reconstructed models dropped slightly. It was probably because the use of ReLU function deactivated some neurons, which is known as the "dying ReLU problem" (21), and the neurons could not be efficiently utilized. Regarding the maximum degree for the SE(3)-Transformers (11), it was beneficial to use up to the degree of 2 features, even although the input and output features utilized only features up to a degree of 1 (scalars and vectors). Fuchs *et al.* observed that there was big improvement when they switched the maximum degree from 1 to 2 (11). We observed a similar trend especially in features for which relationships between other residues were important such as atomic clashes or side chain angle accuracies. On the other hand, Ramachandran angles and rotamer outlier ratios were not affected as much since they could be predicted well with only localized information. Presumably, the use of higher degree hidden features provided inter-residue information, and this resulted in better predictions.

Finally, we examined if more training data, larger model, and secondary structure-dependent rigid body blocks (SS-dep blocks) could improve the performance. When we increased the training dataset size from 5,690 (6k) to 28,914 (29k) with the baseline model, there was marginal improvement. Presumably, the baseline model did not have enough capacity for learning with the bigger training data. Larger models with more layers showed comparable results to the baseline model with the smaller training data set. (**Fig. S10**) However, the performance with smaller models dropped significantly. We observed that training of models with eight linear layer blocks was unstable and occasionally resulted in poor performance. When larger models were trained using the bigger training dataset, the bigger data could be learned by larger models as they had enough capacity to learn them. They outperformed in terms of clash score and side chain torsion accuracies. Similarly, the use of larger crops (384 residues) slightly improved the MolProbity score. The use of SS-dep blocks contributed to accurately model backbone bonded geometries including bond lengths, angles, and Ramachandran angles. The best performance was achieved by aggregating all these components.

***Structure refinement against cryo-EM density map via the cg2all network***

The cg2all network enabled local optimization at a CG representation using scoring functions at both the CG and atomistic representations. (**Algorithm S1**) In the algorithm, an atomistic structure is generated from a CG structure via a cg2all network for the CG representation. An objective function can be defined as a function of both atomistic and the CG representation. Once the objective function is evaluated, the score is backpropagated to get derivatives of the CG structure. Then, the CG structure is updated using the derivative. We applied this algorithm to optimize incorrect protein model structures against cryo-EM density maps as an example usage. Nine protein structures were arbitrarily selected from the test set, and their biological assemblies were set as target structures: 1kq1 (22) (369 residues, 6-mer), 1vim (23) (760 residues, 4-mer), 1g2o (24) (786 residues, 3-mer), 2ibp (25) (814 residues, 2-mer), 1a2z (26) (880 residues, 4-mer), 1s57 (27) (906 residues, 6-mer), 3isr (28) (1,149 residues, 4-mer), 1j0h (29) (1,176 residues, 2-mer), and 1wur (30) (1,848 residues, 10-mer). For those experimental structures, synthetic electron density maps were generated using "molmap" command in UCSF Chimera (31) that employs EMAN2's "pdb2mrc" program (32) at resolutions of 3, 4, 5, 6, 8, and 10 Å. Initial models for local optimization against the electron density maps were predicted by AlphaFold-Multimer (33) with multiple sequence alignments from the ColabFold API (34) and without structural templates using ESMFold (35). We tested our local optimization protocol that utilized an objective function based on both CG and atomistic representations and compared with alternative protocols, including local optimizations at either CG or atomistic representations and molecular dynamics flexible fitting (MDFF) protocol (36).

The objective function for local optimization against electron density map consisted of four objective functions in either atomistic or the CG representation. The first one was the electron density map potential taken from MDFF. (**Eq. 10**)

$$U(\{\vec{\boldsymbol{x}}\}) = \sum_i w_i \max\left(1, 1 - \frac{\Phi(\vec{x}_i) - \Phi_{\mathrm{thr}}}{\Phi_{\mathrm{max}} - \Phi_{\mathrm{thr}}}\right) \tag{10}$$

For this test, we set $\Phi_{\mathrm{thr}}$ to zero and $w_i$ to corresponding atom's atomic mass. The potential was evaluated at the atomistic representation using predicted coordinates from a Cα-trace using the cg2all model network. The second one was backbone bonded potential at the atomistic representation (**Eq. 4**) The third one was a simple CG potential that evaluated pseudo-bond length and angle potential energies (**Eq. 11**) and soft-core van der Waals potential energy (**Eq. 12**).

$$U_{\mathrm{bonded}} = \left(\frac{b_{\mathrm{C\alpha-C\alpha}} - \overline{b_{\mathrm{C\alpha-C\alpha}}}}{\sigma(b_{\mathrm{C\alpha-C\alpha}})}\right)^2 + \left(\frac{\theta_{\mathrm{C\alpha-C\alpha-C\alpha}} - \overline{\theta_{\mathrm{C\alpha-C\alpha-C\alpha}}}}{\sigma(\theta_{\mathrm{C\alpha-C\alpha-C\alpha}})}\right)^2 \tag{11}$$

$$U_{vdW} = \sum_{ij}\left(\min\left(0, d_{ij} - d_{ij,\mathrm{min}}\right)\right)^2 \tag{12}$$

Parameters for the potential functions were obtained from a statistical analysis on the Top8000 structure. As the final one, Cα-Cα distance restraints were applied to residue pairs for which the distances in the initial model were closer than 10 Å. For this experiment, we set relative weights to 1:1:0.1:100.

Initial protein model structures were locally optimized at either residue center-of-mass or Cα-trace representations using a cg2all network-based optimization algorithm (**Algorithm S1**). For this test, we superposed the initial structures onto their target experimental structures using MM-align (37) as initial fits to electron density maps, and the superposition was iteratively updated as well by optimizing the overall structural translation and rotation against the density maps. A structure was optimized using the Adam optimizer (38) for 1,000 or 2,500 steps for AlphaFold and ESMFold models, respectively, and intermediate snapshots were recorded for every 100 steps. The learning rate was updated every step using a cosine annealing scheduler (39), which changed the learning rate from 0.005 to 0.0005 for 200 steps. From the snapshots, a structure with the highest cross-correlation coefficients (CCC) to the target density map was selected as an optimized structure. As alternatives, we performed local optimization with a CG representation using only CG-level objective functions. For the electron density map potential, we used the total mass of a residue as $w_i$ instead. Optimization at a CG representation was carried out in the same way. Then, atomistic structures were generated from Cα-traces using the cg2all network, and the highest CCC structure was selected as an optimized structure using the CG representation. For the local optimization protocol at atomistic resolution, we took the initial minimization step of MDFF protocol implemented by CHARMM-GUI (40) and modified it not to use positional restraints. It performed local energy minimization for 1,000 steps in vacuum using the CHARMM36m force field using NAMD (41), the MDFF electron density map potential, and restraints for secondary structure elements, chirality, and for fixing cis-peptide bonds. We also applied the local optimization protocol at atomistic representation to optimized structures from cg2all network-based optimization and optimization at the CG representations for better agreement of solvent exposed sidechains. As the final option, we carried out the full MDFF protocol implemented by CHARMM-GUI (40).

**Algorithm S1**. Local optimization via a cg2all network

**def** local_optimization_via_network($\{\vec{r}\}$, $N_{step}$ = 1,000, $lr$ = 0.005):

### Initializes global transformation (translation vector and rotation matrix).

$(R, \vec{t}) = (I, \vec{0})$

### Initializes an optimizer for protein structure at a CG representation, $\{\vec{r}\}$
### and the global translation vector and rotation matrix.

optimizer = Optimizer([$\{\vec{r}\}$, $R$, $\vec{t}$], lr=$lr$)

scheduler = CosineAnnealingLR(optimizer, $T_{max}$=200, $\eta_{min}$=0.0005)

### Predicts an atomistic structure using a cg2all network and apply the global transformation

$\{\vec{x}\} = \Theta(\{\vec{r}\})$

$\vec{x}_{center} = \text{mean}(\{\vec{x}\})$

$\{\vec{x}'\} = R \circ (\{\vec{x}\} - \vec{x}_{center}) + \vec{x}_{center} + \vec{t}$

### Local optimization

**for all** $s \in [1, ..., N_{step}]$ **do**

### Evaluates objective function, which is a function of coordinates
### at either an atomistic or the CG representation

score = objective_function($\{\vec{r}\}$, $\{\vec{x}'\}$)

### Backpropagates via the cg2all network, $\Theta$

score.backward()

### Performs a single optimization step that updates coordinates
### at the CG representation and the global transformation

optimizer.step()

optimizer.zero_grad()

scheduler.step()

### Predicts an atomistic structure using updated coordinates and the global transformation

$\{\vec{x}\} = \Theta(\{\vec{r}\})$

$\vec{x}_{center} = \text{mean}(\{\vec{x}\})$

$\{\vec{x}'\} = R \circ (\{\vec{x}\} - \vec{x}_{center}) + \vec{x}_{center} + \vec{t}$

**end for**

**return** $\{\vec{r}\}$, $\{\vec{x}'\}$

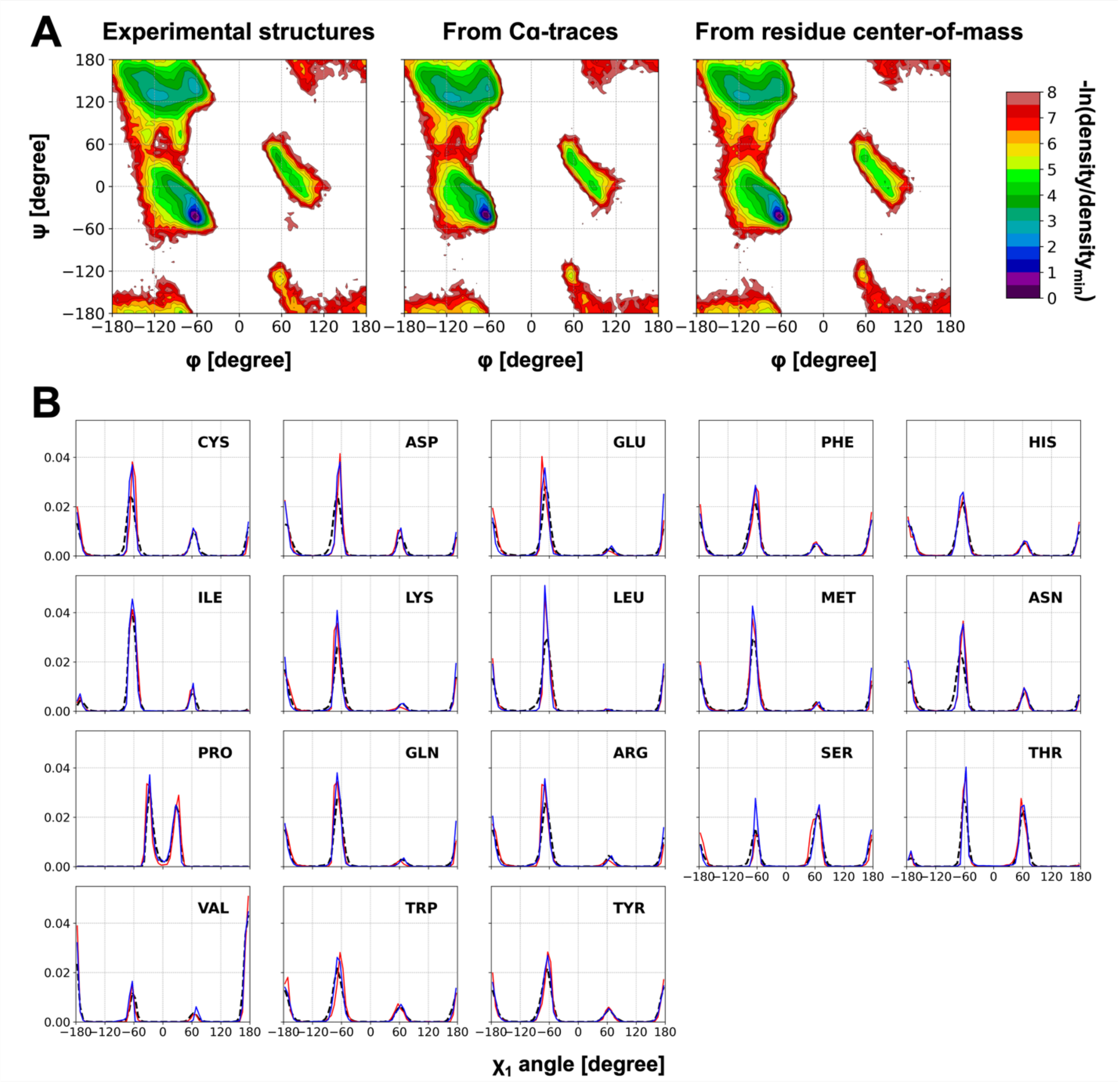


**Figure S1.** Torsion angle distributions of reconstructed all-atom structures from CG structures. Distributions of the Ramachandran angles (A) and the side chain $\chi_1$ angle for each residue (B) are compared between experimental structures and their reconstructions from CG structures. In panel B, data from experimental structures are shown in black dashed lines, while distributions from reconstructed all-atom structures from Cα-traces and residue center-of-mass models are shown as red and blue solid lines, respectively.

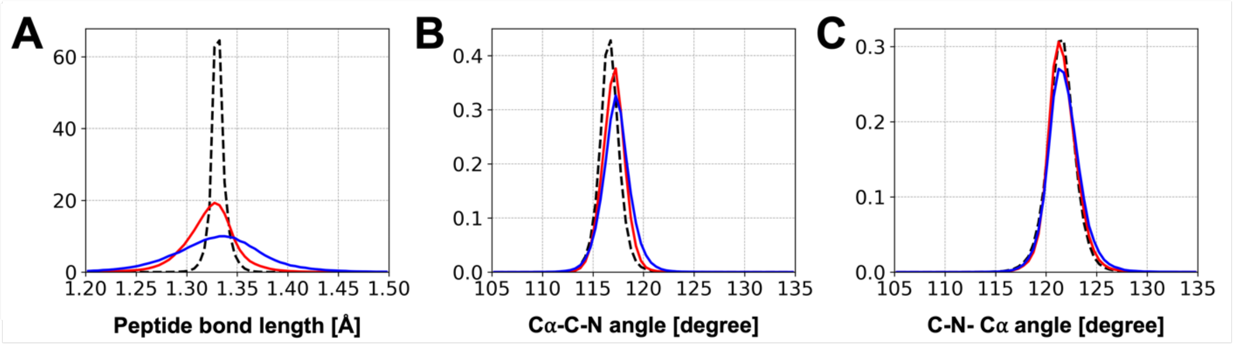


**Figure S2.** Distributions of the peptide bond length between residues (A) and backbone angles containing peptide bonds (B and C). Data from experimental structures are shown in black dashed lines, while distributions from reconstructed all-atom structures from Cα-traces and residue center-of-mass models are shown as red and blue solid lines, respectively.

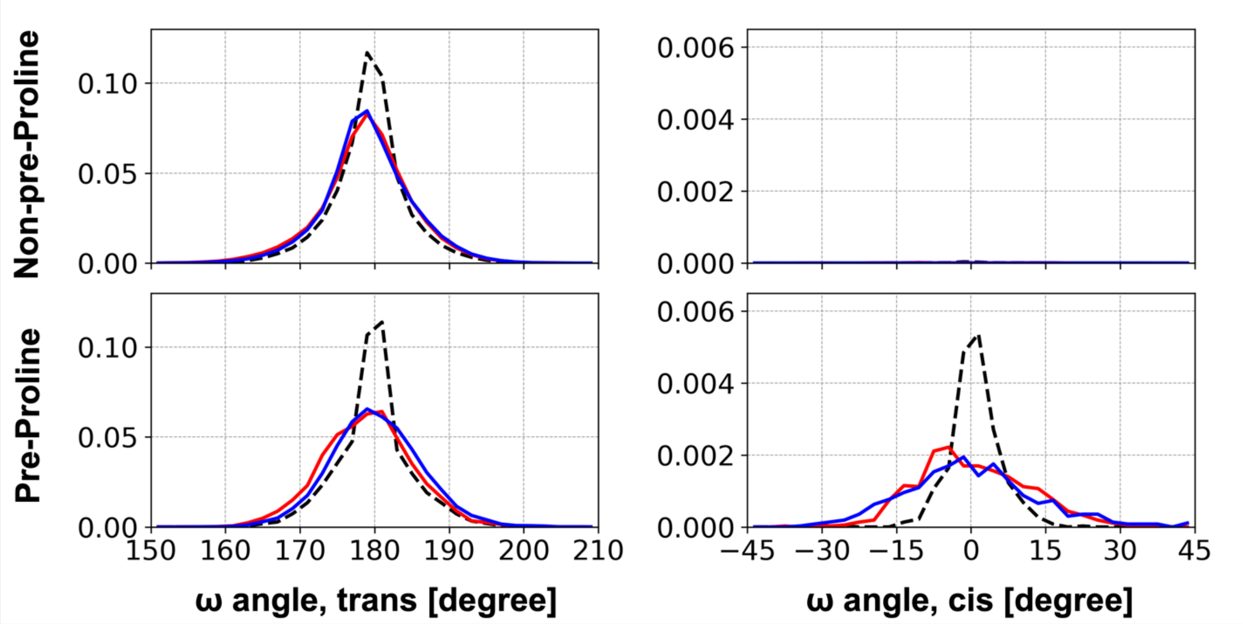


**Figure S3.** Comparisons of omega angle distributions. Data from experimental structures are shown in black dashed lines, while distributions from reconstructed all-atom structures from Cα-traces and residue center-of-mass models are shown as red and blue solid lines, respectively. Comparisons are made separately for non-pre-proline residues (top) and pre-proline residues (bottom) of their trans- (left) and cis- (right) peptide conformations.

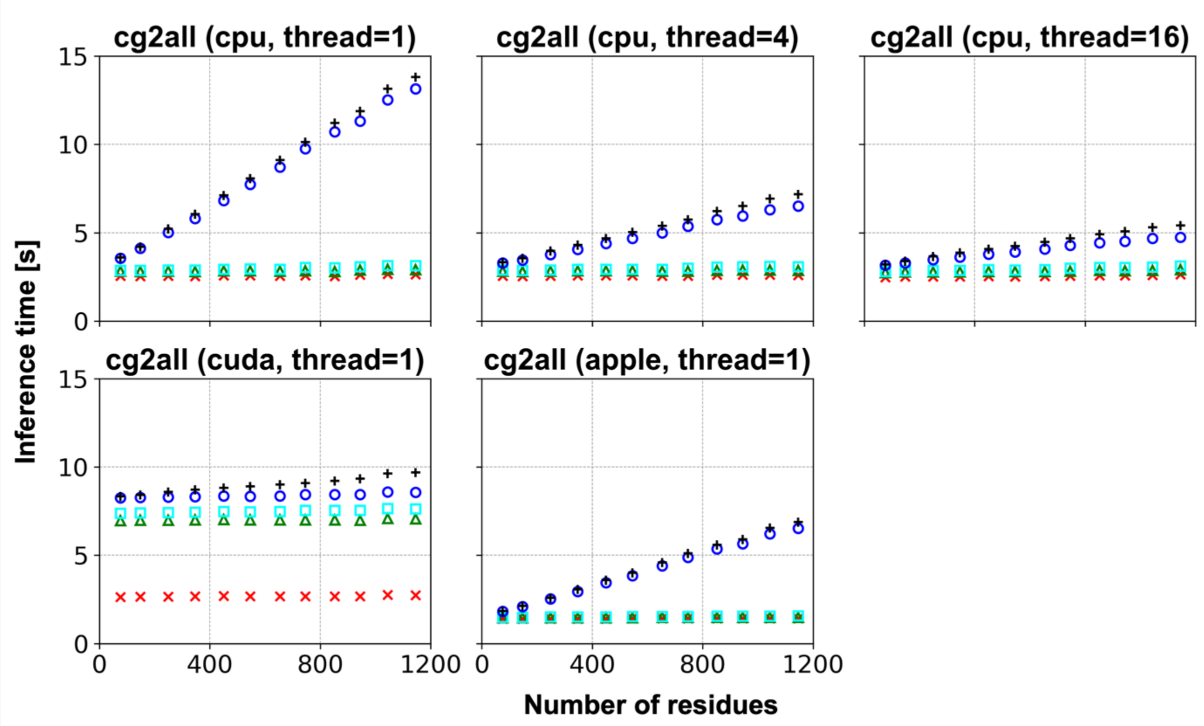


**Figure S4.** Inference time for a conversion from a Cα-trace to an all-atom model as a function of protein size on various devices. Cumulative times from the beginning to the step of loading libraries (red Xs), loading the PyTorch model (green triangles), reading an input PDB file and its pre-processing (cyan squares), forward-pass through the network (blue circles), and writing an output PDB file (black '+') are shown. The "cpu" and "cuda" ran on a Linux system with two Intel Xeon Silver 4214 CPUs (2.20 GHz) (for the "cpu" runs), eight NVIDIA Geforce RTX 2080 Tis (11 GB VRAM) (for the "cuda" runs), and a network file system (NFS) for storage. The "apple" timings were run on an Apple MacBook Pro with an M1 Pro chip (8-core CPU, 14-core GPU, and 16 GB RAM). The "mps" accelerator of PyTorch was not used for the "apple" runs.

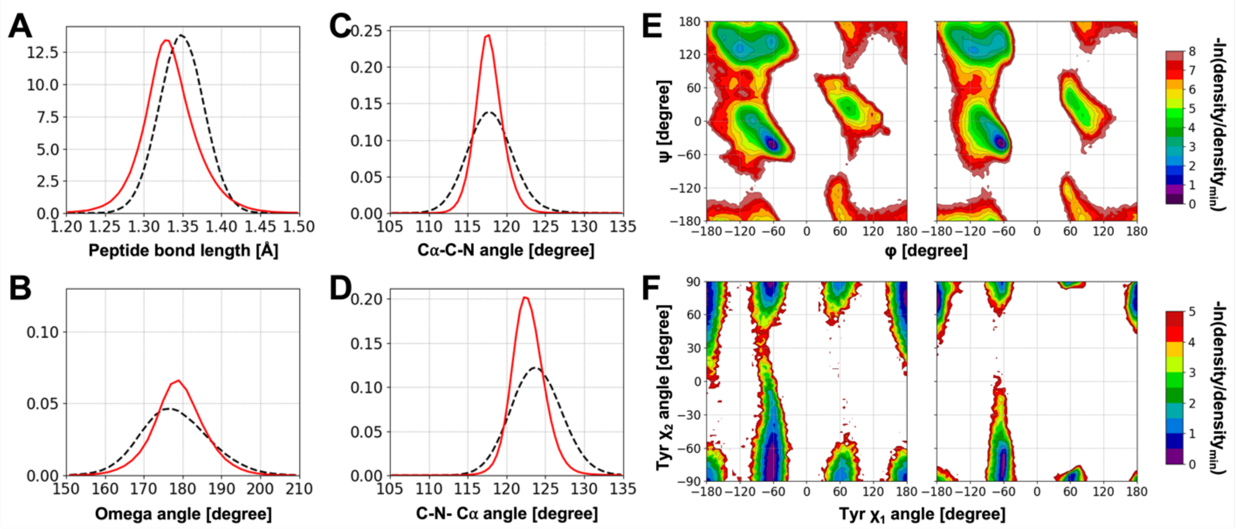


**Figure S5.** Bonded geometry distributions of reconstructed all-atom structures from Cα-traces of all-atom MD simulation snapshots. See **Figures S1-S3** for details.

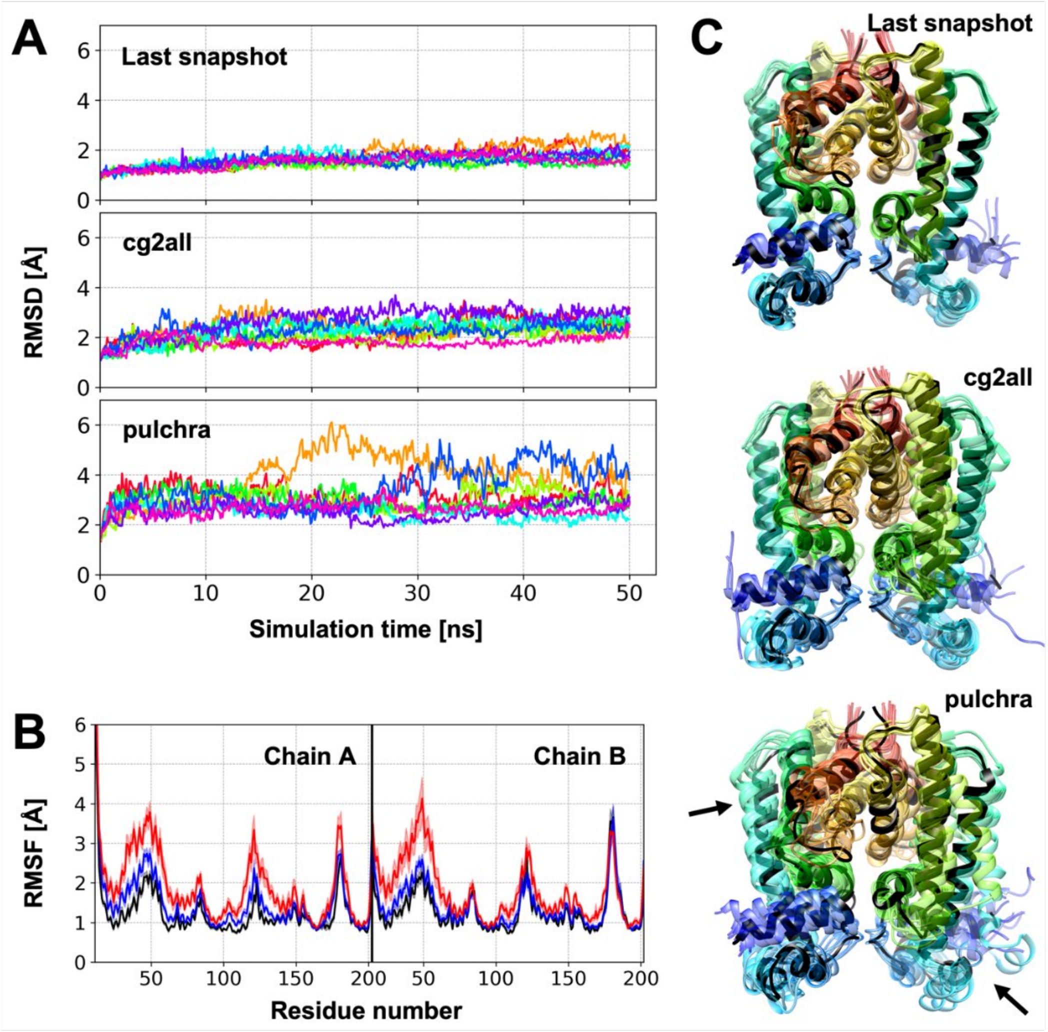


**Figure S6.** The stability of continued all-atom MD simulations starting from reconstructed all-atom models. The last snapshot of a dimeric protein (PDB ID: 2ibd) all-atom simulation was locally minimized using COCOMO model, and the minimized Cα-trace was converted to an all-atom model using cg2all and PULCHRA. Then, eight replicas of all-atom simulations were performed starting from the reconstructed all-atom models after an equilibration step. (A) Cα-RMSD trajectories with respect to their starting model. Each trajectory is colored differently. (B) Residue-wise root-mean-square-fluctuation (RMSF) for the last snapshot, cg2all, and pulchra model are shown in black, blue, and red lines. Standard errors of the value are shown with transparent shades. For the RMSF evaluation, the first 10 ns was discarded as the equilibration process. (C) Ensemble of structures after 10 ns of all-atom simulations (transparent rainbow colors) are compared with their starting structure (black). Highly deviated regions in the PULCHRA simulations are indicated by black arrows.

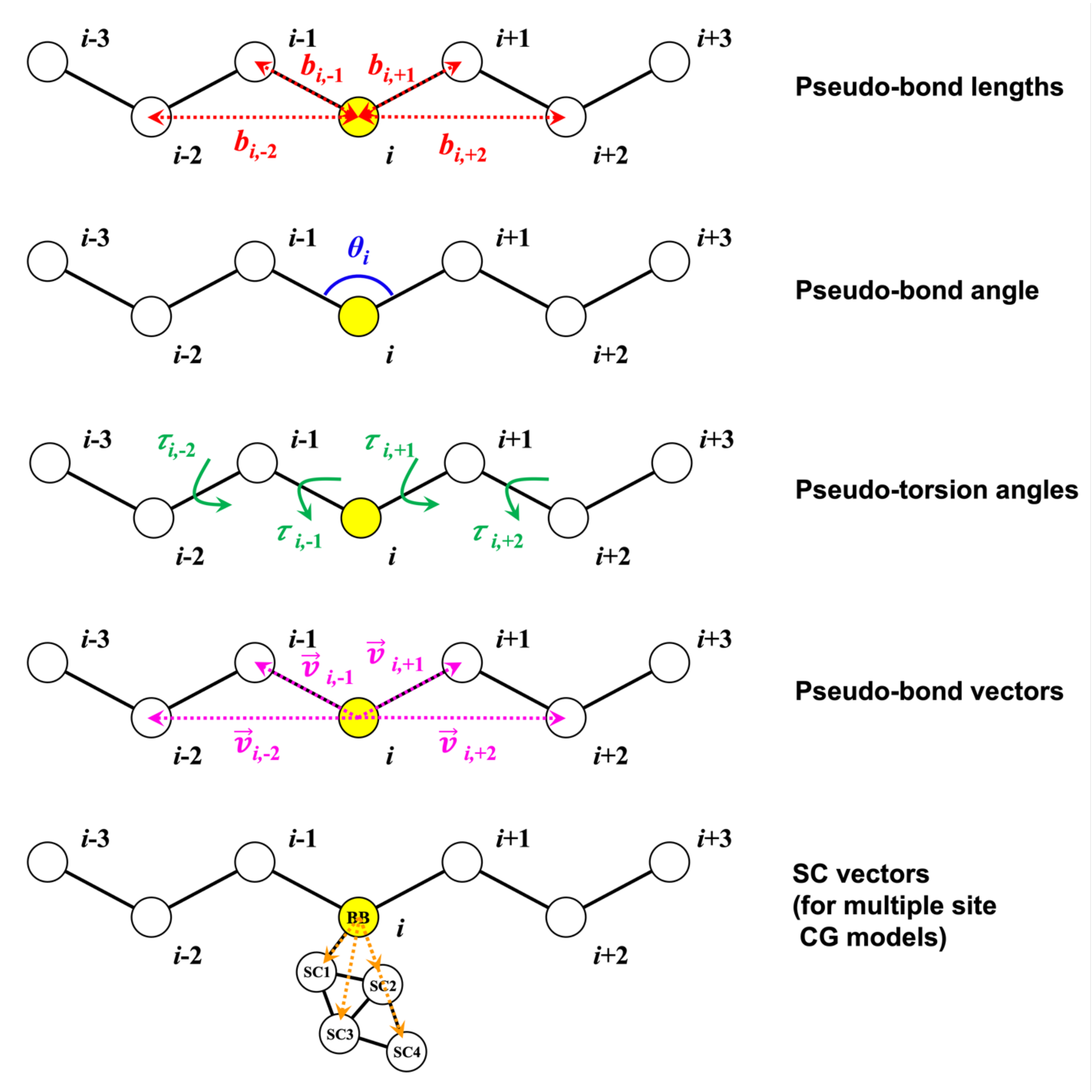


**Figure S7.** Input features of the model. For each residue *i* (yellow circle), pseudo-bond lengths, -bond angle, -torsion angles, -bond vectors were used. For multiple site CG models, sidechain vectors were utilized additionally.

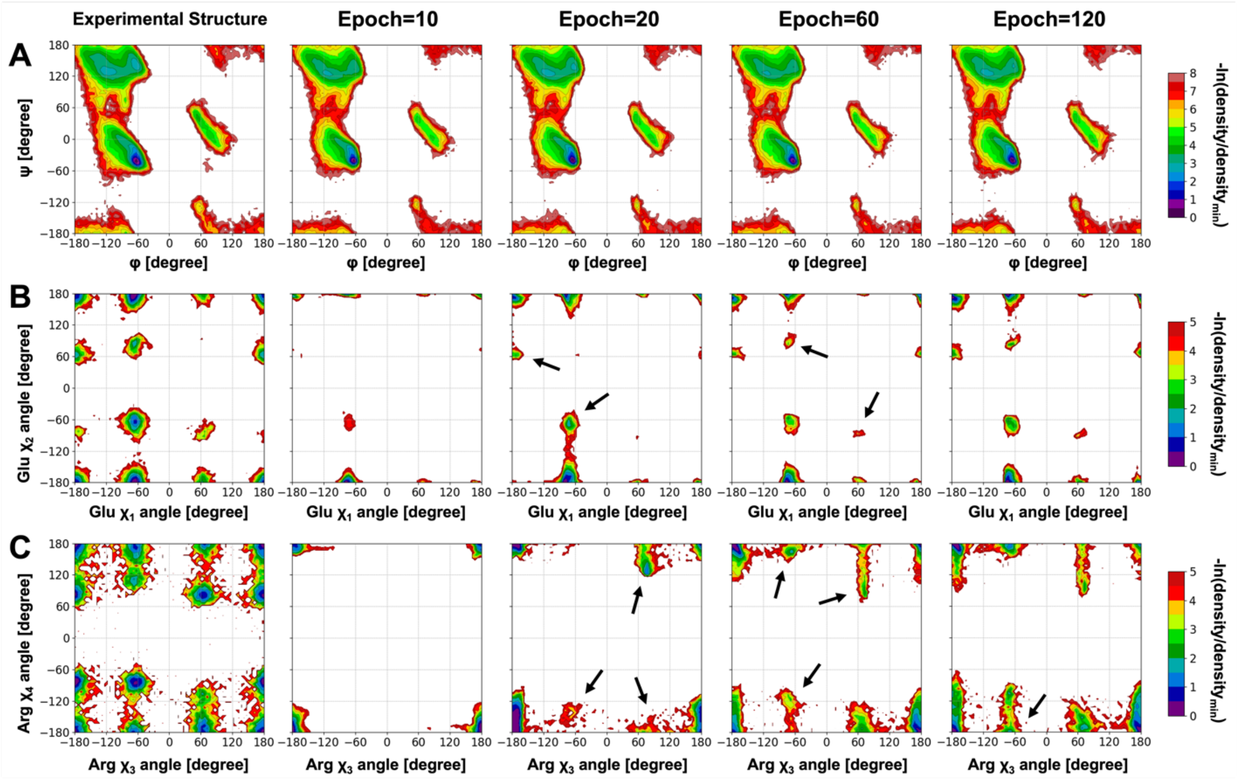


**Figure S8.** Progress in learning structural features with the Cα-based model. Two-dimensional distributions of (A) Ramachandran angles, (B) Glutamate $\chi_1$ and $\chi_2$ angles, and (C) Arginine $\chi_3$ and $\chi_4$ angles at a range of epochs of training are compared with those from the experimental structures. Newly emerging densities compared to a previous step are indicated by black arrows.

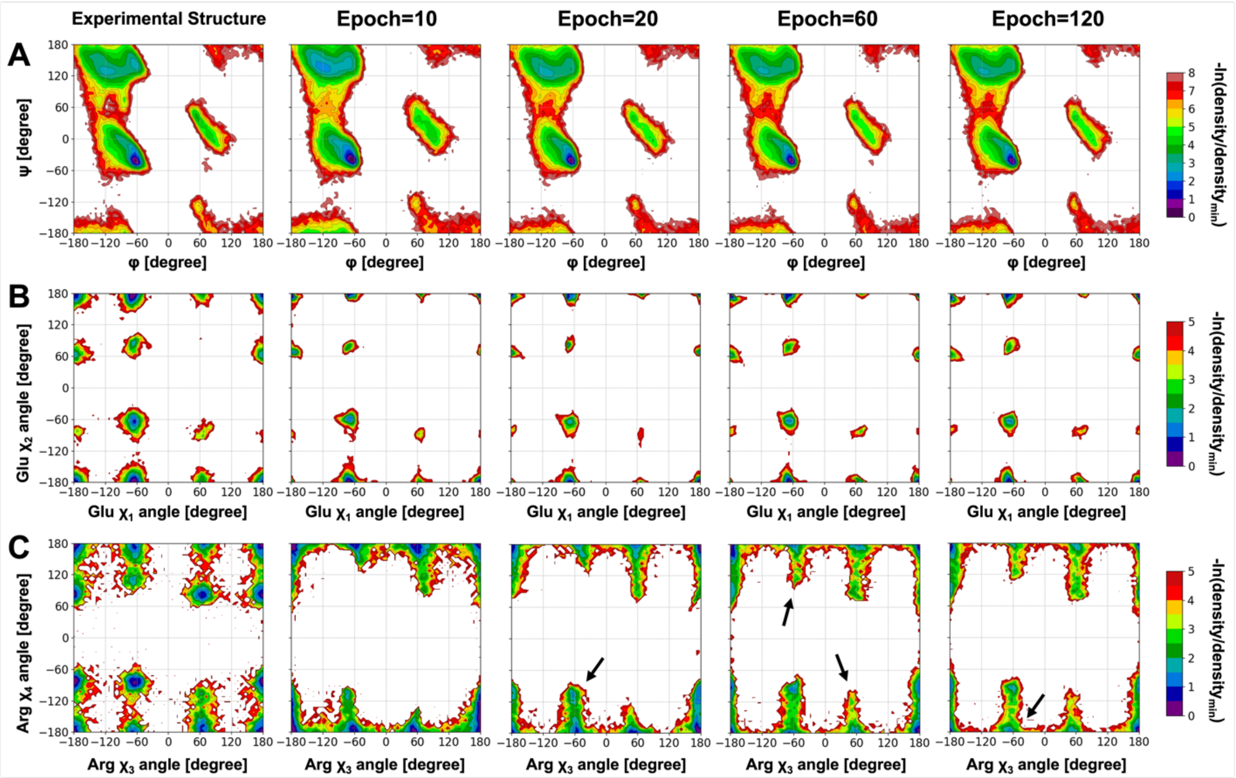


**Figure S9.** Progress in learning structural features with the residue center-of-mass model. See **Fig. S8** for details.

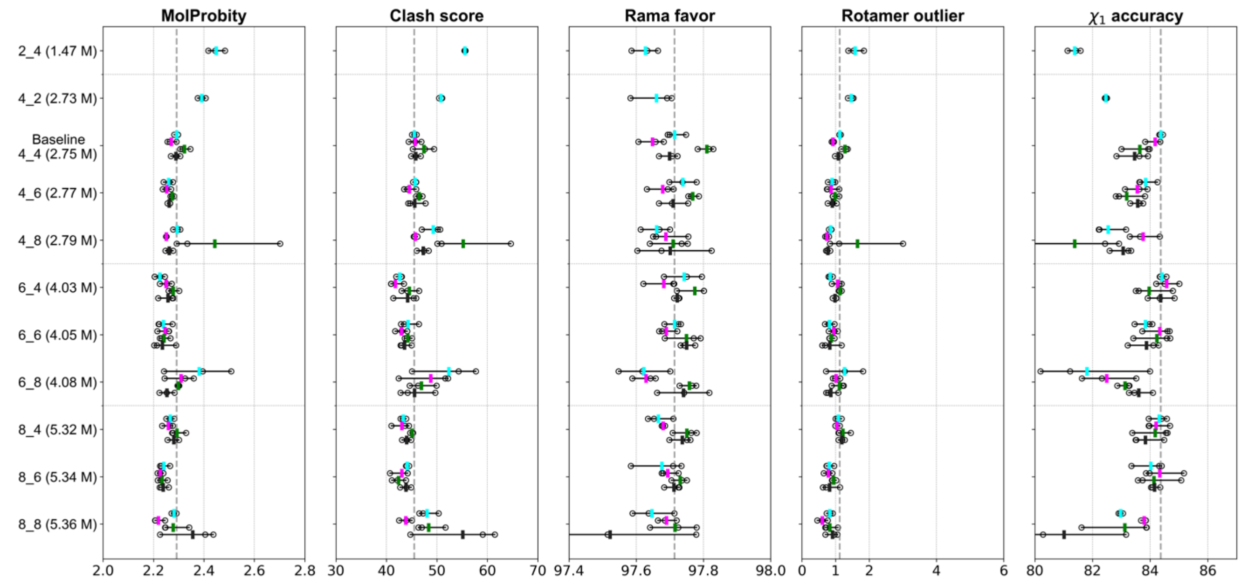


**Figure S10.** Performance as a function of the model size. The numbers of linear layer blocks of the initialization/structure modules (N in **Fig. 1**) and SE(3)-Transformer blocks of the interaction module (M in **Fig. 1**) were varied, and each model is named as N_M in the figure. The total number of parameters are shown in the parentheses. Models were trained with a combination of two different crop size (256 and 384) and the use of secondary structure-dependent rigid body blocks (with and without SS-dep blocks) using the PDB 6k training set. Each of them was trained three times independently, and their results are shown as black circles. The average value of the results is shown as cyan (crop size=256 and without SS-dep blocks), magenta (crop size=384 and without SS-dep blocks), green (crop size=256 and with SS-dep blocks), and dark grey (crop size=384 and with SS-dep blocks) bars. The average value of the baseline model with a crop size of 256 and without SS-dep blocks is also shown as grey dashed lines.

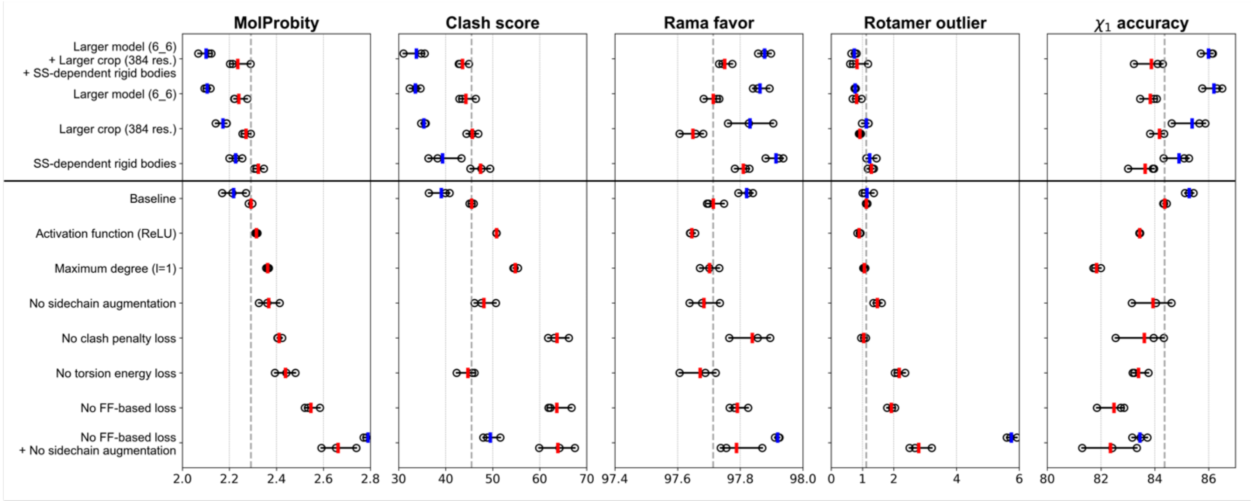


**Figure S11.** Ablation study results using two training sets in various metrics. Each model was trained three times independently, and their results are shown as black circles. The average value of the results is shown as red and blue bars for the training sets of PDB 6k and 29k, respectively. The average value of the baseline model trained with the PDB 6k set is also shown as grey dashed lines.

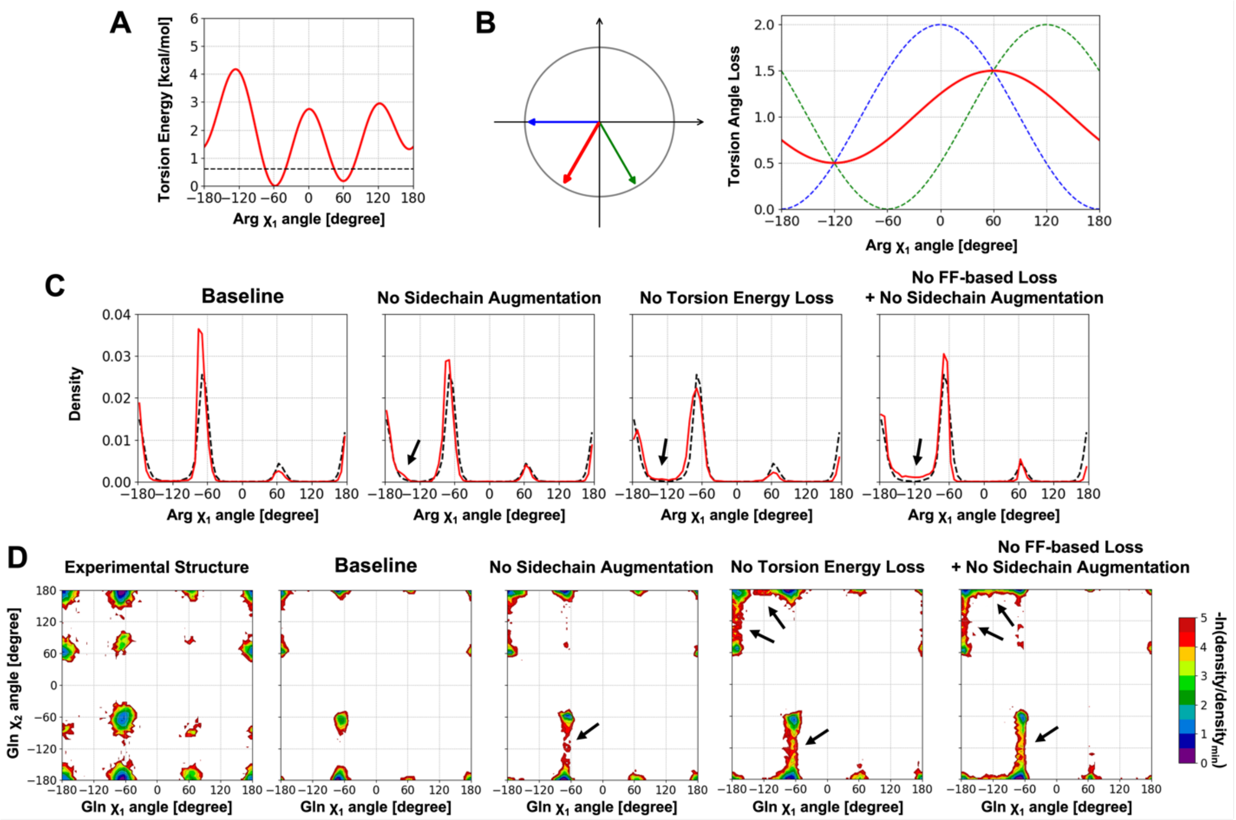


**Figure S12.** Contributions of two side chain-related loss functions. (A) Torsion angle energy profile for Arg $\chi_1$ angle. The torsion energy clamp value ($E_{clamp}$ = 0.6 kcal/mol) is shown as a dashed line. (B) Schematic explanation of the need of sidechain augmentation. Training with two Arg $\chi_1$ angles (-180 and -60 degrees, shown in blue and green) with very similar input features could be resulted in their averaged value, -120 degree (red) because the torsion angle loss function has a minimum at the value. (C) Comparisons of Arginine $\chi_1$ angle distribution between that from the experimental structures (black dashed line) and that from the reconstructed structures from Cα-traces (red solid line). (D) Comparisons of two-dimensional distributions of Glutamine $\chi_1$ and $\chi_2$ angle distribution. (B and C) PDB 29k training set was used for the training of each model, and the other parameters beside the loss function were kept the same. Remaining problematic regions are indicated by black arrows.

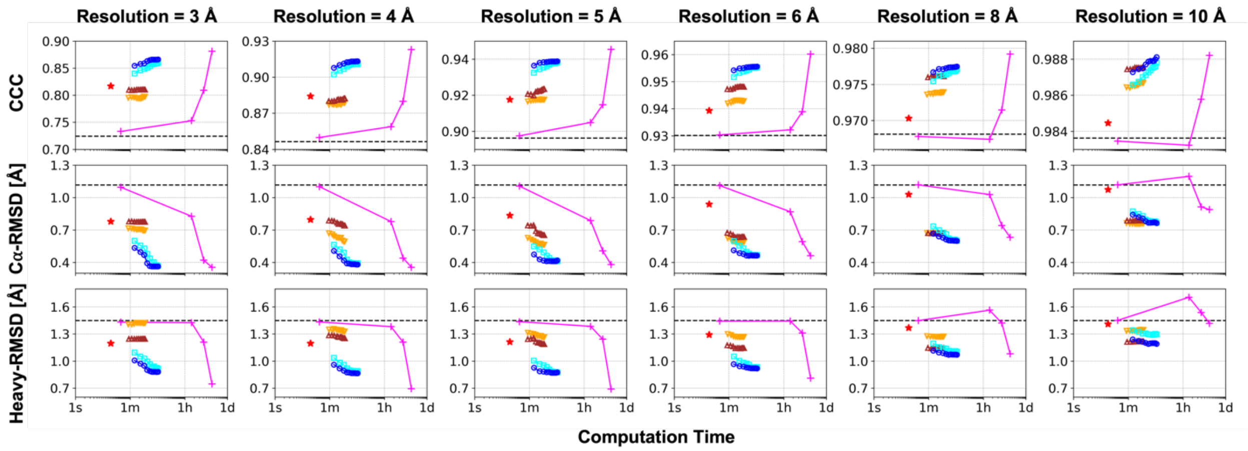


**Figure S13.** Performance of refinement of AlphaFold2 models against cryo-EM density maps. See **Figure 3** for details.

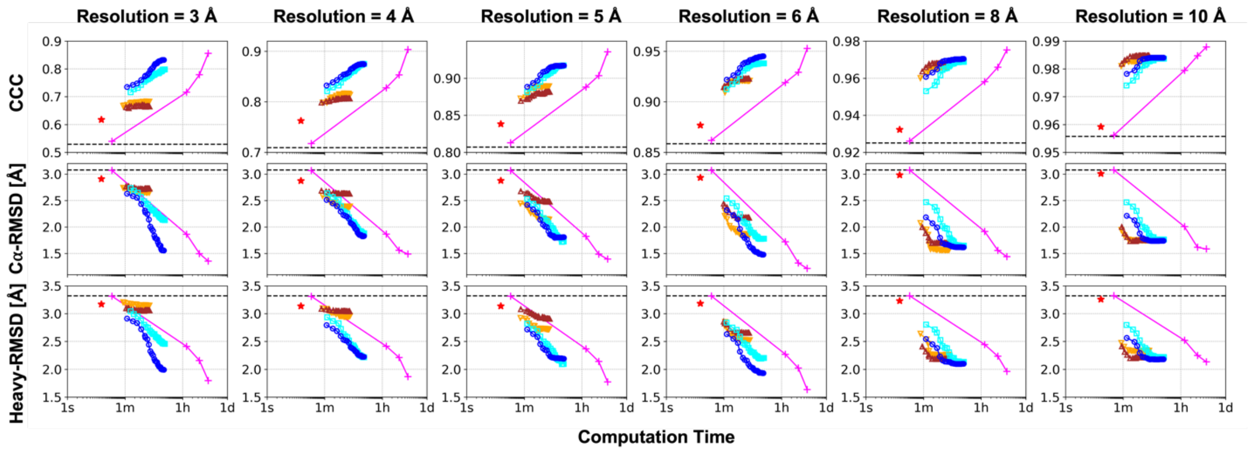


**Figure S14.** Performance of refinement of ESMFold models against cryo-EM density maps. See **Figure 3** for details.

**Table S1.** Performance of conversion to all-atom structures from Cα-traces of simulation snapshots

| Input | Method | RMSD | | χ-angle accuracy[1] | | MolProbity | | | |
|---|---|---|---|---|---|---|---|---|---|
| | | Backbone [Å] | Heavy atom [Å] | $\chi_1$ [%] | $\chi_{1+2}$ [%] | Score | Clash score | Rama favor [%] | Rotamer outlier [%] |
| Cα from all-atom MD snapshots | MD snapshots | | | | | 1.45 | 0.8 | 93.5 | 2.7 |
| | cg2all | **0.25** | **1.18** | **77.7** | **58.8** | 2.31 | 37.1 | **96.4** | 1.0 |
| | w/SCRWL[2] | | 1.27 | 76.2 | 58.2 | **2.20** | **32.5** | **96.4** | **0.0** |
| | PULCHRA[3] | 0.52 | 1.70 | 54.2 | 32.1 | 3.84 | 162.7 | 85.1 | 5.2 |
| | w/SCWRL[2] | | 1.51 | 68.0 | 49.7 | 2.93 | 67.1 | 85.1 | 0.1 |
| | REMO[3.4] | 0.95 | 2.21 | 44.6 | 26.4 | 4.42 | 197.5 | 75.1 | 15.6 |
| | w/SCWRL[2] | | 1.95 | 63.5 | 45.6 | 3.22 | 102.4 | 75.1 | 0.2 |
| Cα after minimization with COCOMO[5] | cg2all | **0.43** | **1.34** | **73.7** | **54.8** | 2.55 | 44.5 | **94.4** | 1.1 |
| | w/SCWRL[2] | | 1.43 | 72.3 | 53.5 | **2.38** | **36.1** | **94.4** | **0.1** |
| | PULCHRA[3] | 0.61 | 1.79 | 52.5 | 31.0 | 3.88 | 156.8 | 83.5 | 5.6 |
| | w/SCWRL[2] | | 1.62 | 65.8 | 47.1 | 2.95 | 65.1 | 83.5 | **0.1** |
| | REMO[3.4] | 1.16 | 2.40 | 43.3 | 25.7 | 4.50 | 194.5 | 69.9 | 17.4 |
| | w/SCWRL[2] | | 2.23 | 59.4 | 41.3 | 3.33 | 116.1 | 69.9 | 0.2 |

[1]Side chain χ-angles were considered accurate when their deviations from experimental values were less than 30 degrees.

[2]Side chains were reconstructed using SCWRL4 after building a backbone structure using other methods (*e.g.*, cg2all, PULCHRA, and REMO).

[3]Chains in multi-chain targets were separately converted to all-atom structures and superposed onto the original Cα-trace.

[4]Conversions of several structures failed because short peptides could not be handled.

[5]All-atom MD simulation snapshots were considered as the ground truth. The average structure change in Cα-RMSD after minimization using COCOMO model was 0.30 Å.

**Video S1.** An example of conversion of simulation trajectory from a Cα-level CG model to the all-atom model as described in **Figure 2D**. A periodic simulation box is shown as a blue box. Cα-traces of proteins are presented as traces on the left side. Their converted all-atom models are shown as cartoon representation on the right side with additional side chain structures as sticks. Each chain has different colors but used the same colors for both representations.
